# Investigating oncoprotein-mediated chromatin dysregulation in *Drosophila melanogaster* uncovers novel modifiers of the developmental impact of H3 K27M and EZHIP

**DOI:** 10.1101/2025.05.26.656136

**Authors:** Samuel D. Krabbenhoft, Tyler E. Masuda, Yadwinder Kaur, Truman J. Do, Siddhant U. Jain, Peter W. Lewis, Melissa M. Harrison

## Abstract

Polycomb Repressive Complex 2 (PRC2) maintains epigenetic repression through the catalysis of H3K27 trimethylation (H3K27me3), which restricts gene expression and preserves developmental gene-regulatory networks. The integrity of PRC2-mediated gene silencing depends critically on the ability of PRC2 to establish and propagate H3K27me3 beyond initial recruitment sites. The oncoproteins EZHIP and histone H3 K27M specifically inhibit this propagation by blocking the allosterically activated state of PRC2, leading to global disruption of H3K27me3 patterns and developmental abnormalities. To uncover chromatin-related pathways intersecting with PRC2 repression, we developed a *Drosophila melanogaster* model with tissue-specific expression of EZHIP and H3 K27M. A targeted RNAi screen of conserved chromatin regulators identified genetic modifiers that when knocked down either enhanced or suppressed developmental phenotypes driven by these PRC2 inhibitors. Strong suppressors, including the Trithorax-group proteins Ash1 and Trx, the PR-DUB complex member Asx, and the nucleoporin Nup153, restored normal development despite persistent depletion of global H3K27me3. Gene expression analyses revealed that suppression reflected reduced expression of genes aberrantly activated following PRC2 inhibition. Together, these findings highlight conserved chromatin-regulatory pathways that intersect with Polycomb to maintain transcriptional balance and support developmental homeostasis.

## Introduction

Polycomb group (PcG) proteins are evolutionarily conserved chromatin regulators that maintain transcriptional repression of developmental genes across cell divisions (Schuettengruber et al. 2017; Laugesen et al. 2019; van Mierlo et al. 2019). Through the coordinated activities of Polycomb Repressive Complex 2 (PRC2), which catalyzes H3K27 trimethylation (H3K27me3), and PRC1, which binds this mark and compacts chromatin, Polycomb complexes establish repressive chromatin domains that preserve lineage fidelity and restrict cellular plasticity (Kingston and Tamkun 2014; Blackledge and Klose 2021). This mode of epigenetic regulation is essential for proper cell fate specification, stem cell maintenance, and the temporal coordination of gene expression during development (Boyer et al. 2006; Pasini et al. 2007; Aloia et al. 2013).

Disruption of Polycomb-mediated repression has been implicated in a wide range of human diseases. In congenital disorders, mutations in PcG components or cofactors give rise to severe developmental syndromes, underscoring their role in embryonic patterning (Schuettengruber et al. 2017; Deevy and Bracken 2019; Laugesen et al. 2019; van Mierlo et al. 2019; Blackledge and Klose 2021). In cancer, aberrant Polycomb activity can drive tumorigenesis through context-dependent mechanisms (Laugesen et al. 2019; Deshmukh et al. 2022; Kim and Kingston 2022). While gain-of-function alterations in PRC2 are observed in multiple malignancies and support oncogenic self-renewal programs, other cancers, particularly a subset of pediatric brain tumors, are characterized by attenuated PRC2 activity (Bender et al. 2013; Lewis et al. 2013; Bayliss et al. 2016; Stafford et al. 2018; Harutyunyan et al. 2019). In diffuse midline gliomas (DMGs) and posterior fossa type A (PFA) ependymomas, this attenuation arises from expression of mutant histone H3 (H3 K27M) or the PRC2-interacting protein EZHIP (CXorf67), both of which result in global depletion of H3K27me3 (Schwartzentruber et al. 2012; Wu et al. 2012; Mohammad et al. 2017; Jain et al. 2019; Piunti et al. 2019).

The H3 K27M and EZHIP oncoproteins do not inhibit PRC2 uniformly across the genome, but instead act as potent, selective inhibitors of its allosterically activated form, which is required for the spreading of H3K27me3 beyond sites of initial recruitment (Chan et al. 2013; Mohammad et al. 2017; Diehl et al. 2019; Harutyunyan et al. 2019; Jain, Rashoff, et al. 2020). By targeting this mechanism, they reduce H3K27me3 genome-wide, with loss at distal regions but retention at sites of initial recruitment. This selective inhibition misdirects, rather than abolishes, PRC2 function and leads to widespread alterations in enhancer activity, transcription-factor occupancy, and chromatin topology in a context-specific manner (Funato et al. 2014; Stafford et al. 2018; Larson et al. 2019). Defining how these perturbations rewire developmental gene-regulatory networks and identifying the pathways that intersect with Polycomb to modulate these effects remain a central challenge.

To investigate how H3 K27M and EZHIP reshape gene-regulatory networks *in vivo*, and to identify pathways that intersect with Polycomb dysfunction, we developed a *Drosophila melanogaster* model for tissue-specific expression of these PRC2 inhibitors. The core machinery of Polycomb repression is deeply conserved in *Drosophila*, which, together with the genetic tractability and cellular resolution of the system, provides a powerful platform for dissecting chromatin regulation in a developmental context (Schwartz and Pirrotta 2007; Simon and Kingston 2013; Kassis et al. 2017). We used this model to compare the phenotypic consequences of H3 K27M and EZHIP expression and performed an RNAi screen targeting conserved chromatin regulators to identify genetic modifiers of PRC2 inhibition. This screen uncovered both enhancers and suppressors of H3 K27M-induced phenotypes. Suppressors restored normal tissue development despite persistent loss of H3K27me3, revealing compensatory mechanisms that limit the consequences of impaired PRC2 spreading. These interactions were conserved across tissues and corresponded with partial reversal of H3 K27M-driven transcriptional changes. Together, our findings demonstrate that selective inhibition of PRC2 spreading disrupts development through a gene-regulatory network that is shared between tissues and reveal conserved pathways that functionally intersect with Polycomb to shape chromatin and transcriptional states *in vivo*.

## Results

### Expression of H3 K27M or EZHIP inhibits allosterically activated PRC2 in *Drosophila*

H3 K27M and EZHIP are potent competitive inhibitors of PRC2, and we previously showed that this inhibition is conserved in *Drosophila* (Jain et al. 2019; Jain, Rashoff, et al. 2020). *In vitro* we demonstrated that EZHIP exhibits greater potency than H3 K27M in inhibiting PRC2 catalytic activity, although the biological consequences of this difference remain poorly understood (Jain et al. 2019). To systematically compare the developmental impact of these two inhibitors, we engineered transgenic *Drosophila* that allowed for expression of either H3 K27M or EZHIP in various tissues. Ubiquitous expression of either inhibitor resulted in lethality, underscoring their deleterious effects on organismal development. Notably, EZHIP expression resulted in lethality at the third-instar larvae stage, whereas H3 K27M expression permitted progression to pupation before death (Figure 1A). Thus, EZHIP expression has more severe developmental consequences than H3 K27M expression. Expression of control transgenes, H3 K27R and EZHIP M406E, harboring amino acid substitutions that do not inhibit PRC2, had no effect on viability. These findings confirm that PRC2 activity is essential for viability in *Drosophila* and establish that the relative potency of H3 K27M and EZHIP in impairing PRC2 function is reflected in their differential developmental consequences *in vivo*.

**Figure 1.**
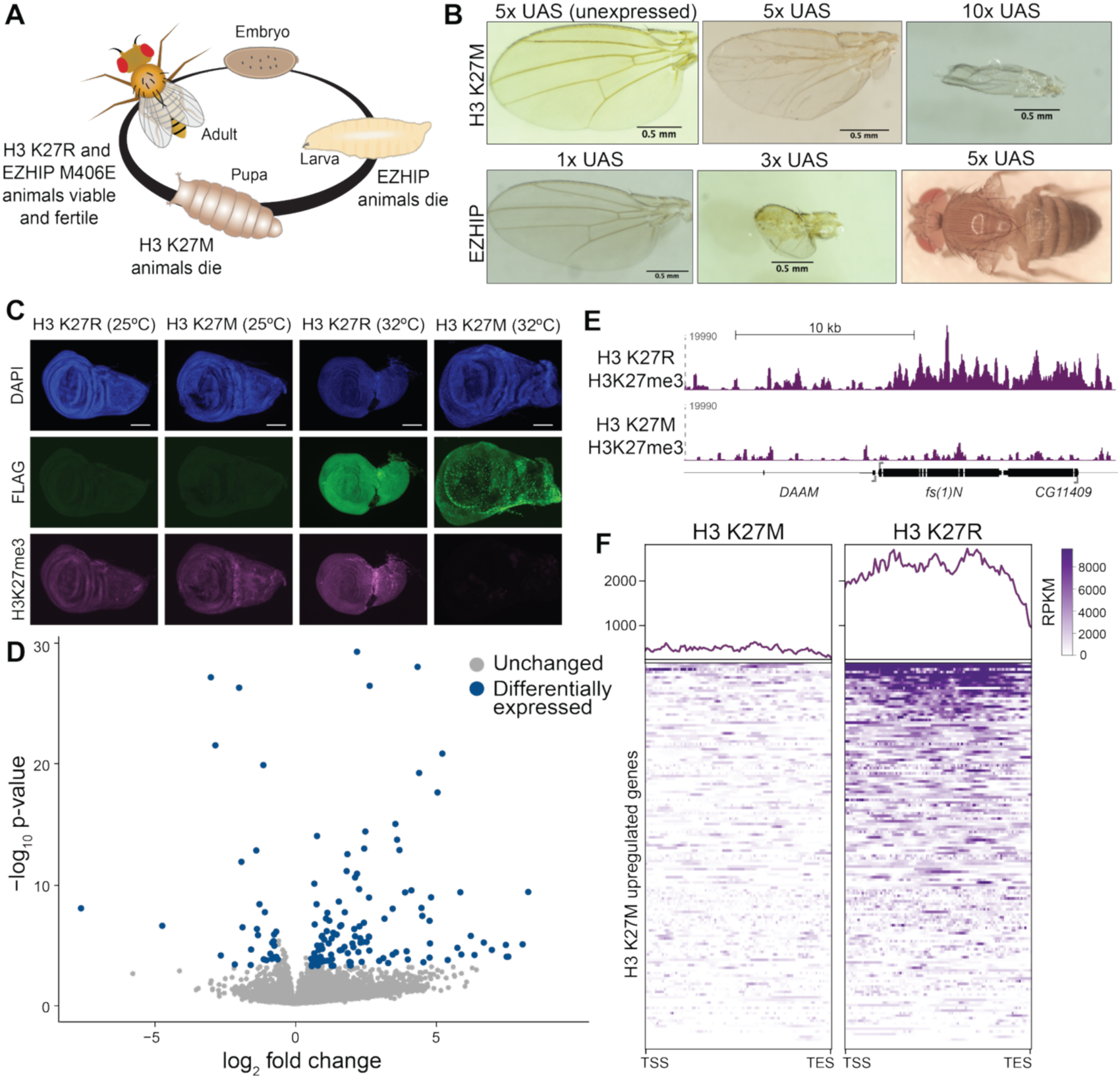
EZHIP and H3K27M expression disrupt PRC2-mediated repression and cause developmental defects. A. Schematic of the *Drosophila* life cycle indicating the developmental stages at which lethality occurs following ubiquitous expression of EZHIP or H3 K27M. B. Adult wing morphology from flies expressing H3 K27M or EZHIP under the control of *nubbin-GAL4*. The number of upstream activating sequences (UAS, GAL4-binding sites) driving transgene expression are indicated above each image. C. Immunostaining of wing discs showing FLAG-tagged H3 K27M or EZHIP (green) or H3K27me3 (magenta). Nuclei are marked by DAPI (blue). Transgenes were expressed for 72 hours. D. Volcano plot showing differential gene expression in wing discs expressing H3 K27M compared to H3 K27R controls. Blue dots indicate upregulated or downregulated genes (adjusted p-value < 0.05, fold changed > 1.5). Gray dots indicate genes with non-significant changes in expression. E. CUT&RUN genome browser tracks showing H3K27me3 signal at *fs(1)N* in wing discs expressing H3 K27M or H3 K27R. *fs(1)N* is upregulated in H3 K27M wing discs as compared to H3 K27R. F. Top: Meta-plots H3K27me3 CUT&RUN signal centered on 148 genes upregulated in H3 K27M-expressing wing discs relative to H3 K27R. Bottom: Heatmaps of spike-in normalized RPKM counts. Genes are sorted by mean H3K27me3 intensity from highest to lowest. TSS, transcription start site. TES, transcription end site.

To directly compare the developmental effects of H3 K27M and EZHIP while avoiding the lethality associated with their ubiquitous expression, we targeted their expression to a dispensable tissue, the wing imaginal disc, using the wing pouch-specific *nub-GAL4* driver. This approach allowed us to assess tissue-intrinsic phenotypes under controlled conditions. Expression of H3 K27M resulted in moderate morphological defects, including wrinkling of the wing blade and disruptions in the vein pattering. By contrast, expression of EZHIP in the same tissue completely abrogated wing development (Figure 1B). These phenotypic differences were sensitive to the level of transgene expression: increasing the number of upstream activating sequences (UAS) driving either transgene led to more severe developmental abnormalities. This dosage-dependent effect is consistent with the known competitive inhibition of PRC2 by both H3 K27M and EZHIP. These observations support the conclusion that the distinct consequences of H3 K27M and EZHIP expression reflect their relative biochemical potency as PRC2 inhibitors and further validate the *Drosophila* wing as a tractable model for dissecting PRC2 regulation *in vivo*.

PRC2 represses gene expression through the catalysis of H3K27me3. To assess the consequences of PRC2 inhibition in the developing wing, we examined both global H3K27me3 levels and gene expression profiles following oncoprotein expression. Because expression of EZHIP caused a near-complete loss of wing tissue, presumably through widespread developmental arrest or altered cell fate, we focused our molecular analysis on the effects of H3 K27M, which produced a less severe and more anatomically intact wing. To uniformly express H3 K27M while maintaining temporal control, we employed the GAL4/GAL80^ts^ system in which the temperature-sensitive GAL80 repressor blocks GAL4-driven H3 K27M expression until animals are shifted to the permissive temperature. This allowed synchronous and ubiquitous induction of H3 K27M across all tissues, including the wing imaginal disc, for a defined 72-hour window (Supplemental Figure 1A) (McGuire et al. 2003). Following induction, immunostaining revealed a marked depletion of H3K27me3 relative to H3 K27R controls, confirming the efficacy of H3 K27M as a PRC2 inhibitor *in vivo* (Figure 1C). To determine consequences of this disruption in chromatin state on gene expression, we performed RNA-seq on dissected wing discs after 72 hours of H3 K27M or H3 K27R expression. We identified 181 differentially expressed genes, of which 148 (82%) were upregulated, consistent with the model in which H3 K27M impairs PRC2-mediated repression of gene expression (Figure 1D, Supplemental Figure 1B, C, Supplemental Table 1,2). Stringent statistical thresholds were applied to define differential expression, requiring both an adjusted p-value < 0.05 and a minimum fold change of 1.5. Gene ontology analysis of the upregulated gene set revealed a significant enrichment for transcripts involved in germline specification, transposon silencing, and piRNA biogenesis, gene categories typically repressed in somatic tissues by Polycomb-mediated mechanisms (Supplemental Figure 1D).

To determine the relationship between changes in gene expression upon H3 K27M expression and the inhibition of PRC2, we performed genome-wide profiling of H3K27me3 occupancy using Cleavage Under Targets and Release Under Nuclease (CUT&RUN). The resulting profiles revealed a genome-wide depletion of H3K27me3 in H3 K27M-expressing discs relative to controls, confirming the broad inhibitory effect of this oncohistone on PRC2 catalytic activity (Figure 1 E, F, Supplemental Figure 2A). Genes upregulated in our RNA-seq dataset showed the most pronounced loss of H3K27me3, linking PRC2 inhibition to transcriptional derepression.

Mechanistically, H3 K27M preferentially inhibits allosterically activated PRC2, which is dispensable for initial genomic recruitment but essential for spreading and maintenance of H3K27me3 domains (Justin et al. 2016; Stafford et al. 2018; Diehl et al. 2019). To test whether this selective inhibition resulted in retention of H3K27me3 at PRC2-recruitment sites, Polycomb Response Elements (PREs), and loss at distal regions, we performed differential CUT&RUN analysis, subtracting the H3K27me3 signal in the H3 K27R-expressing discs from that of the H3 K27M-expressing discs. Consistent with the model of impaired spreading, we identified a global depletion of H3K27me3 signal across the genome with H3K27me3 retained at PREs (Supplemental Figure 2A, B). The pattern of H3K27me3 retention at PREs and depletion elsewhere mirrors observations made in mammalian and *Drosophila* cell culture (Jain, Rashoff, et al. 2020), confirming the functional observation of PRC2 allosteric regulation and its inhibition by H3 K27M in a developing organism. Taken together, our CUT&RUN and RNA-seq data demonstrate that the developmental phenotypes caused by H3 K27M expression arise from impaired spreading of H3K27me3, leading to selective activation of genes that are normally repressed by Polycomb.

### Conserved chromatin-related proteins mediate the H3 K27M wing phenotype

To investigate the mechanisms underlying the H3 K27M-mediated phenotypes, we leveraged our *Drosophila* wing model as a genetically tractable tissue in which to identify additional chromatin-associated pathways that functionally intersect with PRC2 inhibition. We designed a focused RNAi-based screen to identify genes whose depletion modified the severity of the phenotype caused by H3 K27M expression in the wing pouch (Figure 2A). Candidate genes were selected based on three criteria: 1) annotated chromatin-related functions based on Gene Ontology classifications; 2) strong conservation between *Drosophila* and humans, as assessed by the DIOPT ortholog prediction tool (Hu et al. 2011); and 3) confirmed expression in cells derived from the wing disc (Stoiber et al. 2016) (Figure 2B). Based on this selection pipeline and the availability of established *Drosophila* strains, we screened 438 genes, for which we obtained 629 independent RNAi fly lines. Each line was crossed to the *nubbin-GAL4* driver to induce expression of the RNAi construct in the wing pouch, either alone or in combination with the *UAS-H3 K27M*. For each RNAi line, we generated two quantitative phenotypic scores: an “RNAi score” based upon the phenotype caused by expression of the *RNAi* construct alone, and a “screen score” reflecting the phenotype observed in combination with H3 K27M expression. Scores were based on a composite of wing size, vein patterning, wrinkling, and tissue integrity, using a scale from 0 (wildtype) to 10 (severely disrupted) (Figure 2C, Supplemental Table 3). Expression of H3 K27M alone produced a characteristic phenotype with moderate vein defects and tissue wrinkling, resulting in a score of 4 (Figure 2D).

**Figure 2.**
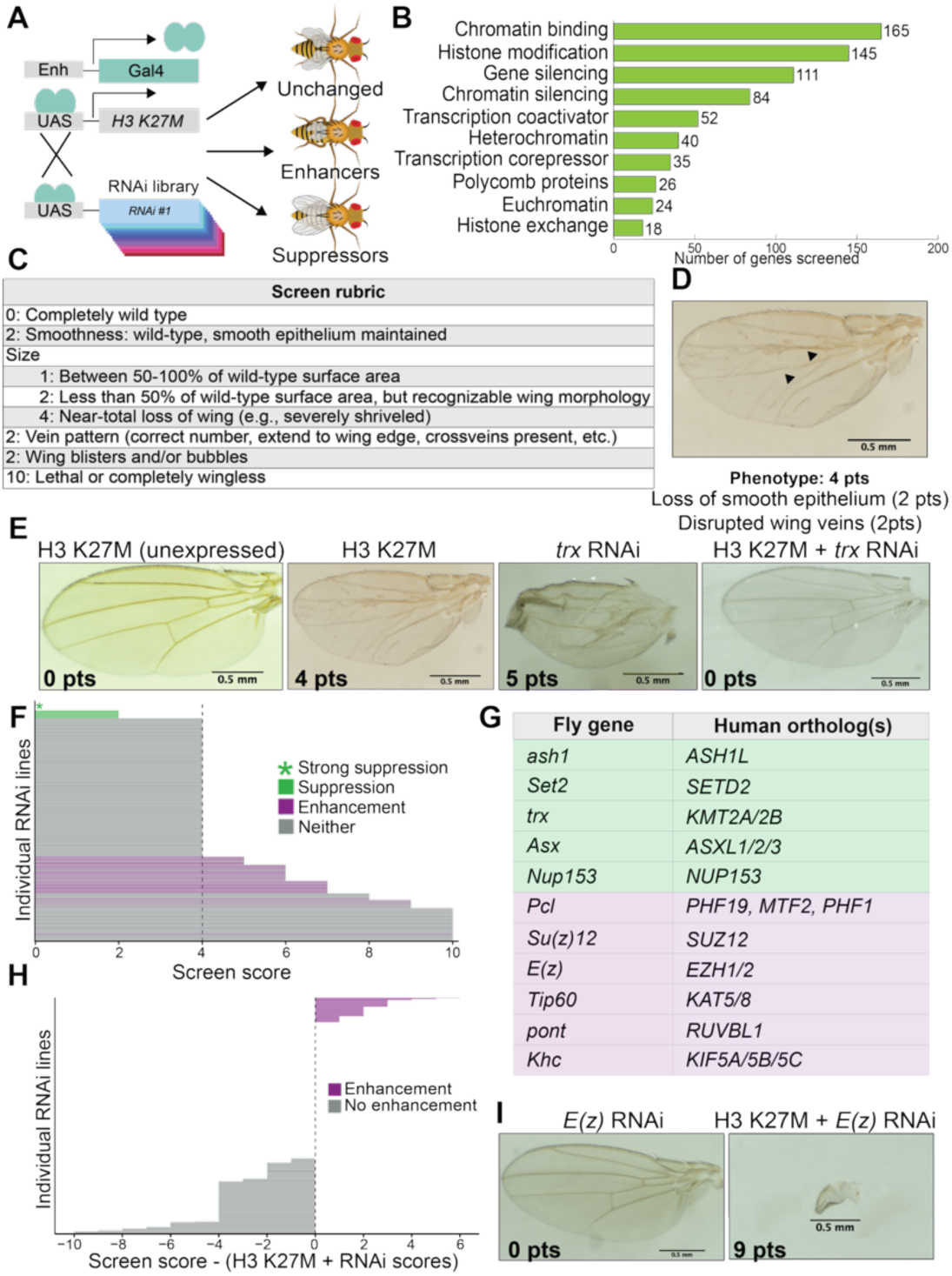
Knockdown of conserved chromatin-associated proteins enhance or suppress the H3 K27M wing phenotype. A. Schematic of the RNAi-based screen to identify chromatin regulators that modify the H3 K27M phenotype. Each RNAi line was expressed alone or co-expressed with H3 K27M in the wing pouch using *nubbin-GAL4*. B. A total of 438 genes encoding proteins with diverse chromatin-related functions were assayed. C. Rubric used to assign phenotypic scores based on wing morphology, incorporating vein patterning, surface texture, and wing size. D. Representative adult wing from a fly expressing H3 K27M. Disrupted vein patterns (arrowheads) and loss of smooth surface are evident. Wing size is largely unaffected. Screen score = 4 pts. E. Representative images showing phenotypic suppression of the H3 K27M wing phenotype by RNAi-mediated knockdown of *trx*. F. Distribution of screen scores for all 630 RNAi lines. Lines with a screen score less than 4 and at least 50% penetrance were classified as suppressors (green); strong suppressors restoring wild-type morphology (score = 0) are indicated with green asterisks. Lines with scores exceeding the sum of the individual H3 K27M and RNAi scores were classified as enhancers (purple). Non-modifying lines are shown in gray. G. Table summarizing all strong suppressors (green) and enhancers (purple) and their human orthologs. H. Identification of enhancers based on screen scores exceeding additive expectations. Enhancers are labeled in purple; non-enhancers in gray. I. Representative image of a strong enhancer demonstrating synergistic phenotypic defects.

Comparative scoring allowed us to identify two categories of modifiers. Suppressors were defined as RNAi lines that reduced the severity of H3 K27M phenotype (screen score < 4), while enhancers exacerbated the phenotype (screen score > additive expectation). We identified 20 genes whose depletion reduced the severity of the H3 K27M phenotype, including five “strong suppressors” that restored a fully wild-type wing morphology (Figure 2E, F, Supplemental Figure 3A, B, Supplemental Table 4). Several suppressors exhibited reciprocal interactions, in which co-expression with H3 K27M mitigated the deleterious effects of the RNAi alone (Figure 2E). Collectively, the strong suppressors are associated with transcriptional activation (Figure 2G).

To identify enhancers, we sought to distinguish genuine synthetic interactions from simple additive or redundant effects. Therefore, we defined enhancers as those genes whose knockdown synergistically exacerbated the developmental defects caused by H3 K27M expression. Specifically, an RNAi line was classified as an enhancer if co-expression with H3 K27M produced a phenotype more severe than the additive effects of expression of H3 K27M and RNAi alone (Figure 2F, H). Using this criterion, we identified 51 enhancers (Supplemental Table 2). Enhancers were divided into “strong” and “weak” subcategories based on severity. Strong enhancers abolished wing development, indicating a profound dependence on the targeted gene for buffering against PRC2 inhibition (Figure 2I, Supplemental Figure 3C).

Functional enrichment analysis revealed that Polycomb proteins were disproportionately represented among the strong enhancers. This included core PRC2 subunits, such as E(z), Su(z)12 and Caf1-55, as well as members of the Polycomb complexes, such as PRC1 and PhoRC (Figure 2G, Supplemental Figure 3D). These findings support the model that H3 K27M does not abolish PRC2 activity entirely, but rather attenuates its spreading function, leaving residual repression at PREs. Knockdown of additional Polycomb components likely further destabilizes this residual activity, resulting in a catastrophic loss of chromatin-mediated repression and severe developmental disruption.

To rigorously test whether the enhancer and suppressor phenotypes were specific to H3 K27M-mediated PRC2 inhibition rather than due to nonspecific effects of histone transgene overexpression or GAL4/UAS dosage, we performed crosses for expression of the H3 K27R control. For each enhancer and suppressor line, we co-expressed the RNAi construct with H3 K27R and assigned an “H3 K27R score.” In nearly all cases, the H3 K27R score matched the RNAi score alone, indicating that the phenotypic modifications observed in combination with H3 K27M were not due to non-specific interactions with histone overexpression (Supplemental Table 3).

### Suppressors restore normal development to tissues with attenuated PRC2 activity

To address how depletion of specific chromatin regulators could counteract the developmental defects induced by H3 K27M, we determined whether suppression was attributable to changes in transgene expression or restoration of PRC2 function. We first determined whether suppression correlated with reduced levels of the H3 K27M transgene, which could trivially explain the amelioration of phenotypes. To address this, we performed immunostaining for the FLAG epitope on H3 K27M either in wing discs dissected from animals expressing H3 K27M alone or in combination with strong suppressors. In all cases, FLAG signal intensity was comparable between suppressed and unsuppressed tissues, indicating that suppression was not due to silencing of transgene expression (Figure 3A and data not shown). We next assessed whether suppression resulted from restoration of PRC2 enzymatic activity and H3K27me3 levels. Despite dramatic phenotypic rescue, H3K27me3 levels remained low in these tissues (Figure 3A). These findings demonstrate that the suppression of the H3 K27M phenotype does not require restoration of H3K27me3 levels. Rather, we conclude that additional chromatin-related proteins act downstream of or in parallel to PRC2 to mediate the transcriptional and developmental consequences of its inhibition by H3 K27M, and that depletion of these factors can compensate for the disrupted Polycomb function.

**Figure 3.**
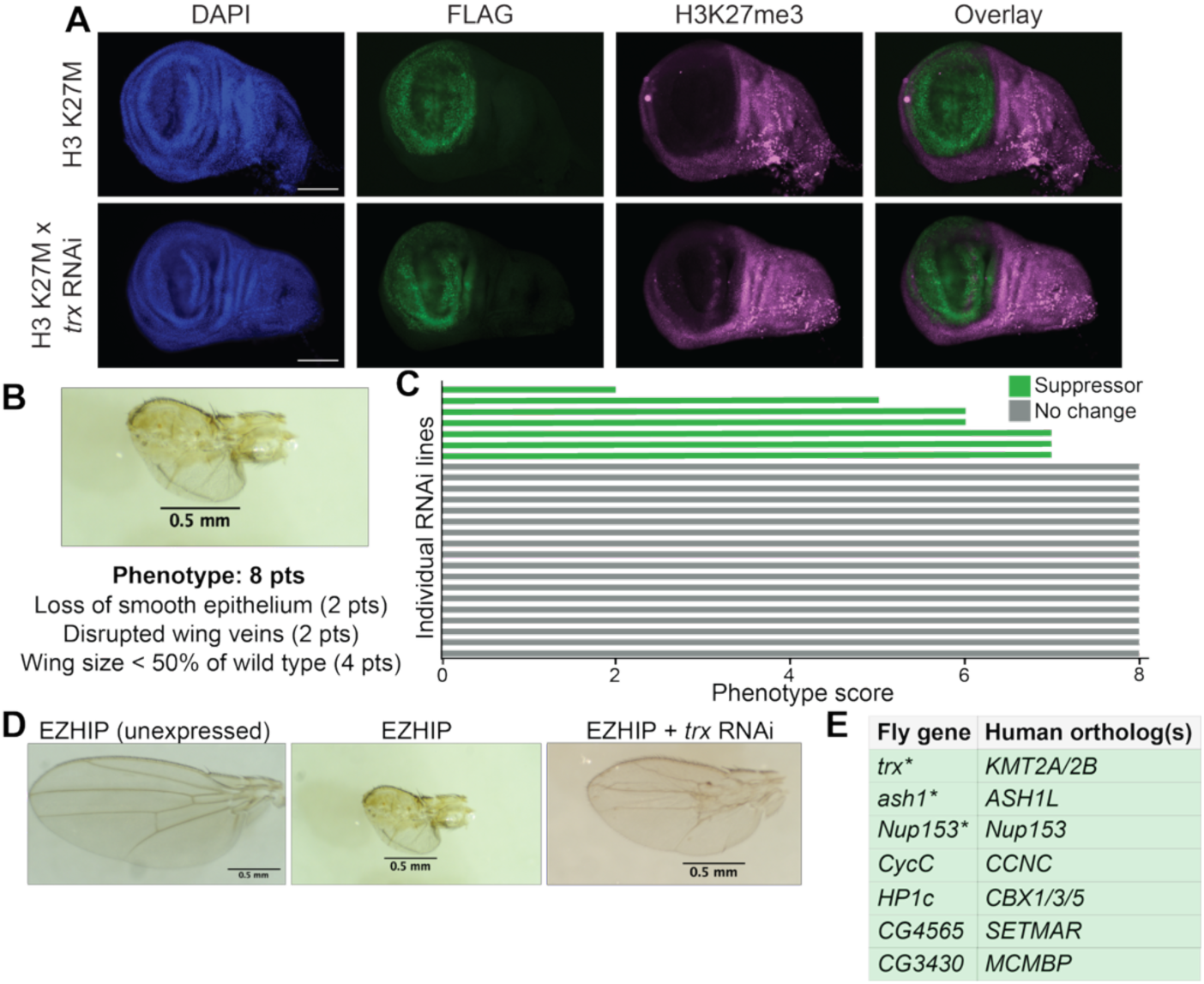
Suppressors restore normal development to tissues with inhibited PRC2 activity. A. Immunostaining of third instar larval wing imaginal discs showing FLAG-tagged H3 K27M (green) and H3K27me3 (magenta). Transgenes expressed by the *nubbin-GAL4* driver. DAPI labels DNA (blue). B. Representative image of the wing phenotype caused by EZHIP expression, annotated with the corresponding quantitative screen score. Screen score = 8 pts. C. Screen scores for all RNAi lines tested in the EZHIP screen. All RNAi lines identified as H3 K27M suppressors were evaluated. Lines with scores less than 8 were classified as EZHIP suppressors (green). Non-suppressors are shown in gray. D. Example images of a suppressor that partially rescues EZHIP-induced wing defects. E. Table of EZHIP suppressors and their human orthologs. Asterisks (*) indicate suppressors that were also classified as strong H3 K27M suppressors.

To assess whether these findings extend to EZHIP, we evaluated whether the same suppressor could rescue phenotypes caused by its overexpression. Like H3 K27M, EZHIP expression in the wing pouch decreases H3K27me3 levels (Supplemental Figure 4A), but with a far more severe developmental outcome - complete loss of wing tissue (Figure 3B). This more extreme phenotype could result from EZHIP being a more potent inhibitor of PRC2, or from additional PRC2-independent effects on development (Jain et al. 2019). To distinguish between these possibilities, we tested whether our identified suppressors and enhancers could influence EZHIP expression phenotypes similar to their effects on H3 K27M-expressing wings. Due to the severity of the EZHIP-associated phenotype, we were unable to reliably score enhancers. We therefore restricted our analysis to the previously identified 20 H3 K27M suppressors. Of these, six suppressors conferred partial restoration of wing development in EZHIP expressing flies (Figures 3C-E, Supplemental Table 3). These included several of the strongest suppressors from the H3 K27M screen. By contrast, RNAi lines that failed to rescue EZHIP-induced defects were classified as weak H3 K27M suppressors, suggesting that the magnitude of rescue may correlate with the severity of the PRC2 inhibition. The partial rescue observed in EZHIP-expressing wings further supports the idea that the underlying mechanism of suppression is not specific to one PRC2 inhibitor but instead reflects a common requirement for additional chromatin regulators to enforce the gene expression consequences of PRC2 dysfunction.

Taken together, these data indicate that the wing phenotypes induced by H3 K27M and EZHIP expression arise from PRC2 inhibition in conjunction with other chromatin-based processes. The ability of suppressor knockdown to restore wing development without reestablishing H3K27me3 highlights a functional interplay between PRC2 and transcription-promoting pathways. These findings suggest that the developmental consequences of impaired PRC2 activity are not immutable and that compensation via modulation of specific chromatin regulators can restore development even in the context of sustained PRC2 inhibition.

### H3 K27M modifiers are robust across multiple tissue types

To determine whether the chromatin regulators that modulate the developmental consequences of H3 K27M expression in the wing are similarly required in other tissues, we extended our analysis to the *Drosophila* eye imaginal disc. Expression of H3 K27M in the eye disc using the *eyeless-GAL4, GMR-GAL4* driver resulted in decreased H3K27me3 levels and disruption of the well-patterned photoreceptor units of the eye, resulting in a rough-eye phenotype (Figure 4A). These phenotypes were qualitatively distinct from those observed in the wing discs but similarly reflected impaired PRC2 function (Supplemental Figure 4B). We tested whether the enhancer or suppressor RNAi lines identified in the screen modified the H3 K27M-induced phenotypes in the eye disc. Strikingly, many of the same suppressors rescued wild-type eye development (Figure 4A, Supplemental Figure 4C). Likewise, enhancers exacerbated the H3 K27M-induced disorganization, causing more severe morphological defects in the eye. Notably, as in the wing, some suppressors impaired eye development when expressed alone but were suppressed by H3 K27M. These observations suggest that the genetic interactions identified in the wing are not tissue-specific but reflect general features of the chromatin landscape that govern PRC2-dependent gene regulation in multiple developmental settings.

**Figure 4.**
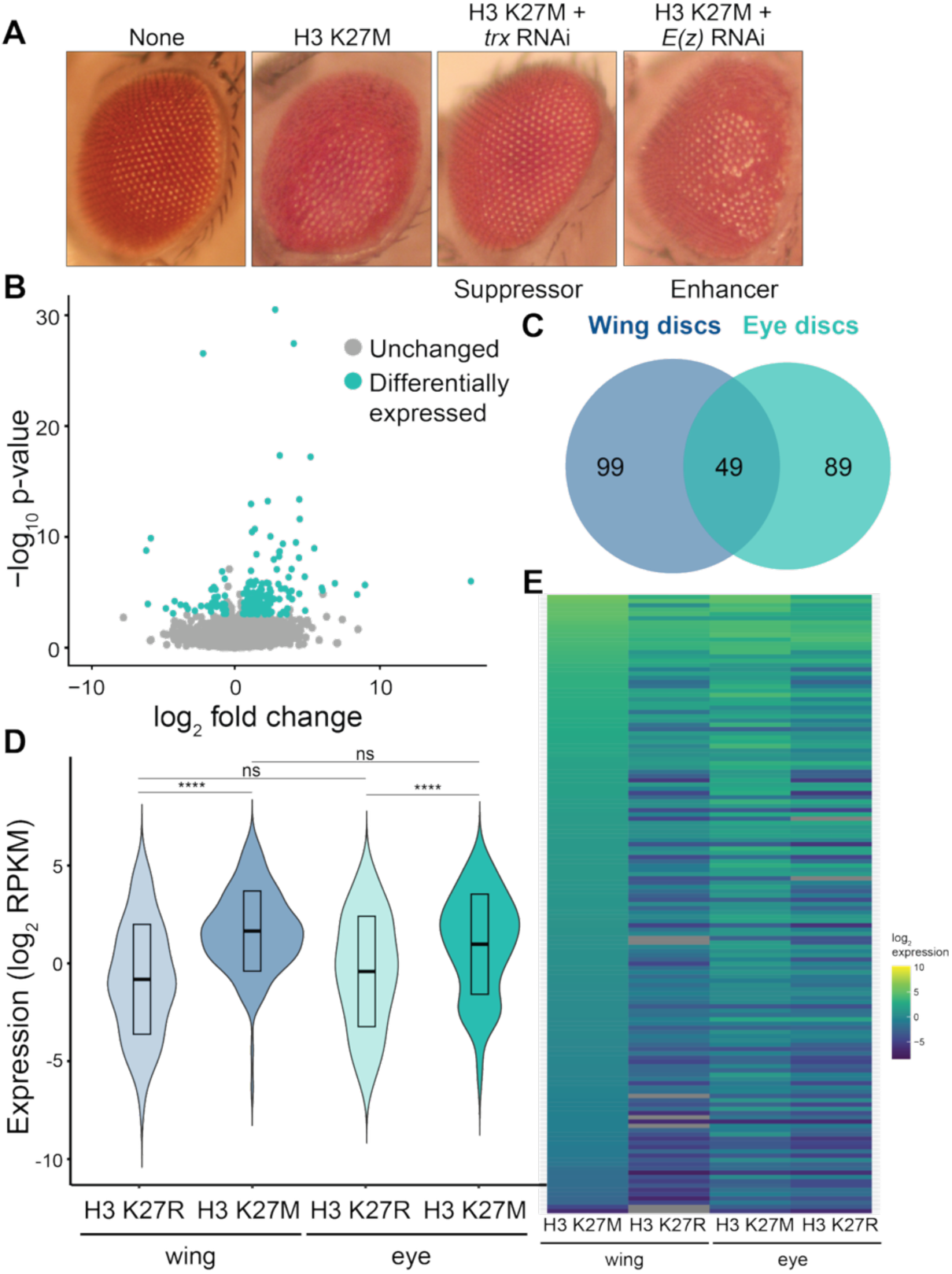
Suppression and enhancement of the H3 K27M phenotype is shared between tissues. A. Representative image of the rough-eye phenotype caused by H3 K27M expression in the eye disc along with an example of suppression and enhancement. Transgenes were expressed using *eyeless-GAL4, GMR-GAL4.* B. Volcano plot showing differential gene expression in eye discs expressing H3 K27M as compared to H3 K27R controls. Blue dots indicate upregulated or downregulated genes (adjusted p-value < 0.05, fold changed > 1.5). Gray dots indicate genes with non-significant changes in expression. C. Venn diagram illustrating the overlap of genes upregulated by H3 K27M in wing and eye discs. D. Violin plots showing the average expression (log_2_ RPKM) of the 148 genes upregulated by H3 K27M in wing discs as compared to H3 K27R controls. Expression is shown for the genotypes and tissues indicated below. ns, not significant. ****, adjusted p-value < 0.0001 (one-way ANOVA). E. Heatmaps of the log_2_ expression of the 148 genes upregulated in wing discs for the genotypes indicated below. Genes are ordered by average expression in H3 K27M wing discs.

To assess whether the phenotypic similarities between the wing and eye tissues extended to the underlying disruption in gene expression, we performed RNA-seq on wing and eye-antennal discs following 72 hours of inducible, continuous H3 K27M or H3 K27R expression. In the eye-antennal disc, H3 K27M expression led to the upregulation of 137 genes and downregulation of 41 genes relative to H3 K27R (Figure 4B, Supplemental Figure 5A). The set of genes upregulated in H3 K27M-expressing eye discs had similar levels of expression in a previously published data set from H3 K27M-expressing eye discs, supporting the reproducibility of our findings (Chaouch et al. 2021) (Supplemental Figure 5B, Supplemental Table 1,2). In contrast to their data using overexpression of H3 as a control, we used H3 K27R, which may account for differences between control samples. To evaluate the degree to which changes in gene expression overlap between tissues, we identified the set of genes upregulated by H3 K27M in both wing and eye discs. Forty-nine genes were shared between the two datasets (Figure 4C), a number significantly greater than expected by chance. To more broadly determine whether H3 K27M affects similar regulatory programs in both tissues, we calculated the average levels of gene expression of the 148 genes upregulated in H3 K27M wing discs as compared to controls. This set of genes was globally upregulated in H3 K27M-expressing eye discs as compared to controls, suggesting that PRC2 represses a common cohort of target genes in both tissues (Figure 4E, F). A reciprocal analysis using the 137 upregulated genes from the eye-antennal disc confirmed these genes also exhibited increased average level of expression in H3 K27M wing discs as compared to H3 K27R controls (Supplemental Figure 5C, D). By contrast, far fewer genes were downregulated by H3 K27M in both tissues, and only six were shared between tissues (Supplemental Figure 5E). Overall, these data support the conclusion that H3 K27M disrupts a conserved set of gene-regulatory networks in multiple tissues, and that the modifiers identified in the wing act through generalizable mechanisms that are operative across developmental contexts.

### Suppressors counteract H3 K27M transcriptional changes

To understand the molecular basis of the phenotypic suppression in animals co-expressing H3 K27M and RNAi against strong suppressors, we asked whether these modifiers act at the level of transcription to counteract the aberrant gene expression profile induced by PRC2 inhibition. Using our temperature-inducible system, we performed RNA-seq on wing discs expressing H3 K27M or H3 K27R with RNAi targeting each of four strong suppressors: *Additional sex combs* (*Asx*), *absent, small, or homeotic discs 1* (*ash1*), *trithorax* (*trx*), and *Nucleoporin 153kD* (*Nup153*). Following 72 hours of transgene induction, we harvested wing discs for transcriptome analysis and assessed differential gene expression across all genotypes (Supplemental Table 1). Each of the four suppressors significantly altered expression of hundreds of genes in H3 K27M wing discs. Strikingly, RNAi of *Asx*, *ash1*, and *trx* preferentially reduced the expression of genes upregulated by H3 K27M expression in wing discs, suggesting that these suppressors are directly required for the ectopic transcriptional activity caused by PRC2 inhibition (Supplemental Figure 6A-C). Importantly, genes downregulated in discs co-expressing H3 K27M and suppressor RNAi as compared to H3 K27M alone were expressed at comparable levels when the same RNAi constructs were combined with H3 K27R, indicating that these expression changes are attributable to the loss of suppressor function itself and not simply additive effects of the transgenes (Supplemental Figure 6A-D).

To directly quantify the extent to which suppressors reversed the transcriptional effects of H3 K27M, we focused on the set of 148 genes previously identified as significantly upregulated by H3 K27M relative to H3 K27R. We categorized whether the expression level of these 148 genes increased, decreased, or remained unchanged when the suppressor RNAi was co-expressed with H3 K27M. Between 22% and 55% of the genes were decreased upon suppressor knockdown, whereas only 3-13% were further upregulated (Figure 5A). When we assessed the average expression level of all 148 genes, we observed that RNAi against *Asx*, *ash1*, and *trx* substantially restored the average expression of these genes to levels similar to the H3 K27R control (Figure 5B, Supplemental Figure 7A). These results demonstrated that the ability of these suppressors to rescue H3 K27M-induced developmental phenotypes is linked to their role in facilitating activation of a subset of PRC2-repressed genes.

**Figure 5.**
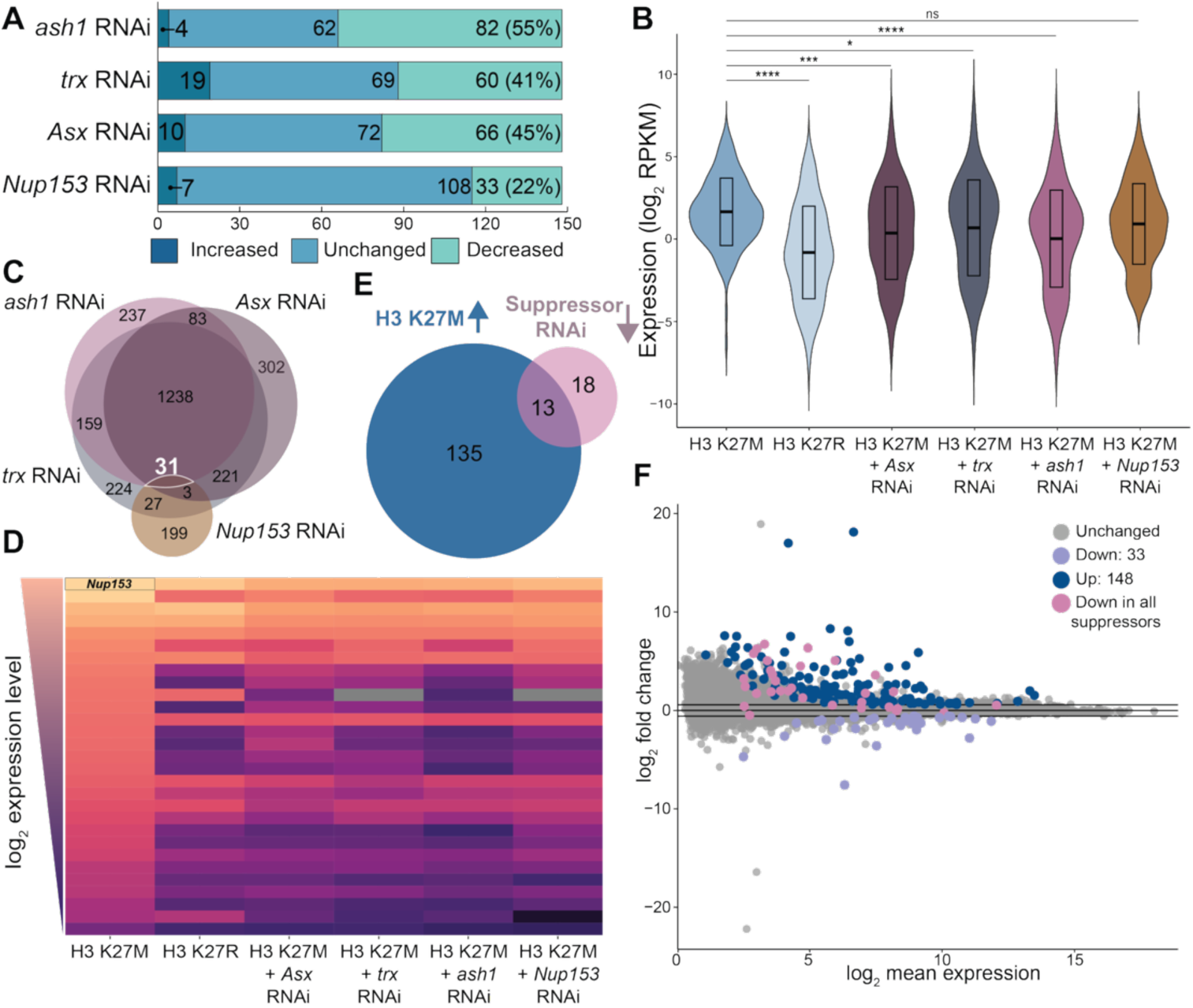
Suppressors counteract H3 K27M-induced transcriptional changes. A. Number of the 148 genes upregulated in H3 K27M-expressing wing discus that are further upregulated, downregulated or unchanged when co-expressed with RNAi targeting the indicated suppressors, compared to H3 K27M alone. B. Average expression (log_2_ RPKM) of the 148 genes upregulated in H3K2M in the genotypes shown below. ns, not significant. *, adjusted p-value < 0.05. ***, adjusted p-value < 0.001. ****, adjusted p-value < 0.0001(one-way ANOVA). C. Venn diagram of the overlap among genes downregulated in wing discs expressing H3 K27M along with RNAi of *ash1, trx, Nup153,* or *Asx1* compared to H3 K27M alone. The subset of 31 genes downregulated by all four suppressors is highlighted. D. Heatmap of the expression (log_2_ RPKM) of the 31 genes downregulated by all four suppressors. *Nup153* was among these genes and is labeled. E. Venn diagram illustrating the overlap of genes downregulated by RNAi of all four suppressors and genes upregulated by H3 K27M expression. F. MA plot showing the differentially expressed genes in wing discs expressing H3 K27M as compared to H3 K27R. Genes upregulated by H3 K27M are shown in dark blue; downregulated genes in lavender. The 31 genes downregulated by all four suppressors are highlighted in pink. Gray dots represent genes with non-significant changes.

Because *Asx, ash1,* and *trx,* all partially rescued the gene expression defects caused by H3 K27M, we tested whether they have broadly similar effects on gene expression. Consistent with their overlapping phenotypic effects, we found that 1,269 genes were significantly downregulated by all three suppressors in H3 K27M wing discs (Figure 4C, Supplemental Figure 7B). This gene set included 45 of the 148 H3 K27M-upregulated genes, reinforcing the hypothesis that these factors are directly required for the increased expression of a subset of PRC2-target loci (Supplemental Figure 7C). Furthermore, many of the same genes were also downregulated when suppressors were expressed in the H3 K27R background, indicating that these proteins control a shared set of target genes (Supplemental Figure 7D).

In contrast to *Asx*, *ash1*, and *trx*, RNAi against *Nup153* did not alter the average expression level of the 148 H3 K27M-upregulated genes, despite its ability to strongly suppress the phenotype caused by H3 K27M expression (Figure 5B). Of the 1,269 genes downregulated by *Asx*, *ash1*, and *trx* RNAi in H3 K27M-expressing discs, only 31 were also downregulated by *Nup153* RNAi, suggesting that Nup153 operates through a largely distinct transcriptional program (Figure 5C-F Supplemental Figure 7E). Despite this limited overlap, nearly half (42%) of the genes that were downregulated by all four suppressors were also part of the H3 K27M-upregulated set (Figure 5F), demonstrating a partial convergence at key target loci. It is noteworthy that levels of *Nup153* itself were downregulated by all four suppressors, hinting at a potential regulatory feedback loop. These findings raise the possibility that suppression of Nup153 may be a critical determinant of phenotypic rescue.

## Discussion

In this study, we developed *Drosophila melanogaster*, the organism in which the Polycomb group proteins were initially discovered, as a model to interrogate the mechanisms by which H3 K27M and EZHIP, two oncoproteins that inhibit allosterically activated PRC2, disrupt normal development. *Drosophila* offers a powerful platform to dissect chromatin-based mechanisms due to the high degree of conservation and robust developmental patterning amenable to phenotypic screening (Yamamoto et al. 2024). By expressing H3 K27M and EZHIP in developing tissues, we modeled the developmental consequences of PRC2 inhibitionc and employed a targeted RNAi screen to identify chromatin-related modifiers that either enhanced or suppressed these phenotypes. Our results revealed that while PRC2 inhibition is necessary to drive developmental defects, it is not sufficient. Suppressor knockdown restored wild-type development despite persistent PRC2 inhibition, suggesting that a delicate equilibrium between repressive chromatin spreading (via H3K27me3) and transcriptional activation is essential for proper gene expression during development (Figure 6).

**Figure 6.**
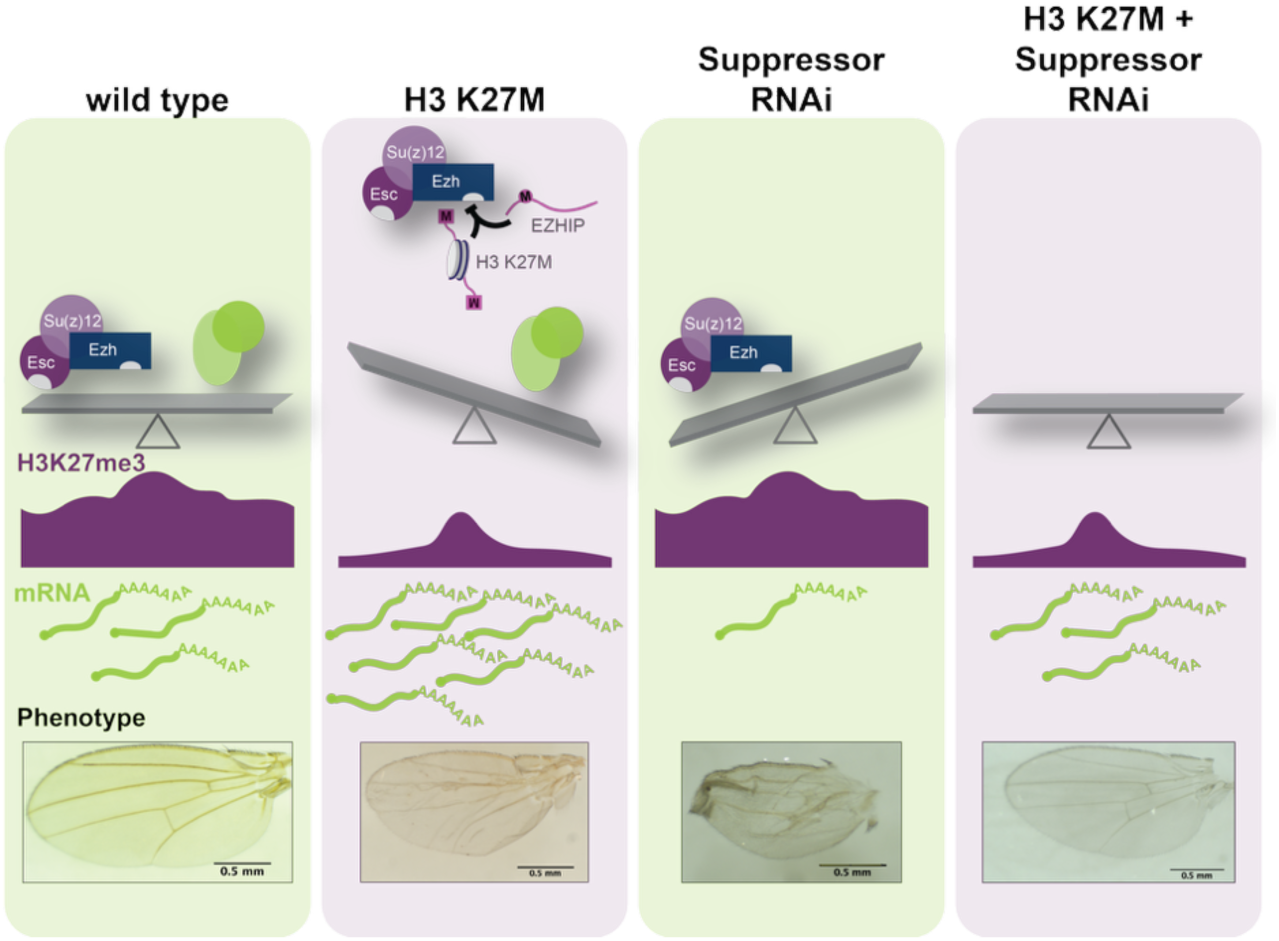
A balance between PRC2-mediated gene silencing and transcriptional activators is essential for normal gene expression and development. During normal development, the interplay between PRC2 activity and the activity of chromatin-associated transcriptional activators, such as Asx, Trx, Ash1, Set2, and Nup153, maintains appropriate gene expression levels, supporting normal wing development (*left*). Inhibition of allosterically activated PRC2 by oncoproteins such as H3 K27M or EZHIP restricts the spreading of H3K27me3, leading to aberrant transcriptional activation and developmental defects (middle panel). Similarly, RNAi-mediated depletion of suppressors alone can disrupt this balance, resulting in reduced gene expression and impaired wing development (middle panel). Strikingly, when PRC2 inhibition is combined with knockdown of suppressors, transcriptional balance is restored and normal development proceeds, despite persistently low global H3K27me3 levels (*right*).

### PRC2 inhibition by H3 K27M or EZHIP results in the developmental defects

By directly comparing EZHIP and H3 K27M, we established that both proteins interfere with PRC2 function in a dose-dependent manner and that the severity of phenotypes scales with oncoprotein expression level. This is consistent with their shared mode of action as competitive inhibitors of PRC2. Biochemically, EZHIP is a more potent inhibitor than H3 K27M (Jain et al. 2019), and our *in vivo* findings support this differential potency: EZHIP expression caused earlier and more severe developmental phenotypes than H3 K27M expression. These phenotypic differences were not observed with control transgenes encoding non-inhibitory mutants (H3 K27R and EZHIP M406E), further confirming that developmental disruption is driven by PRC2 inhibition rather than secondary effects of transgene expression.

Importantly, our screen was specifically designed to identify genetic modifiers that influence the developmental consequences of inhibiting the allosterically activated form of PRC2. Using tissue-specific drivers to avoid the lethality caused by ubiquitous expression and by focusing on phenotypes rather than cell death, we expanded the phenotypic landscape beyond that typically assessed in cell culture-based synthetic lethal screens (Kumar et al. 2017; Berlandi et al. 2019; Yu et al. 2021; Panditharatna et al. 2022; Zhang et al. 2022; Gannon et al. 2023). This enabled the discovery of conserved chromatin regulatory pathways that collaborate with or buffer against PRC2-mediated repression *in vivo*.

### Oncoprotein phenotypes are mediated by conserved chromatin modifiers across tissues

H3 K27M and EZHIP are oncogenic in the developing human hindbrain, suggesting a potential tissue-specific susceptibility (Monje et al. 2011; Mendez et al. 2020). To explore whether the chromatin modifiers identified in our screen act in a tissue-restricted manner, we tested their ability to modify H3 K27M phenotypes in two *Drosophila* tissues. We observed similar enhancer and suppressor interactions in both wing and eye-antennal discs, as well as a shared transcriptional response to H3 K27M across tissues. These findings imply that the underlying regulatory pathways modulated by PRC2 inhibition are broadly conserved and not restricted to a single developmental lineage. The ability of specific chromatin modifiers to restore gene expression and development in both tissues reinforces the conclusion that these regulators are core mediators of PRC2-dependent gene regulation.

### Residual PRC2 activity contributes to developmental outcomes in H3 K27M-expressing tissues

Our genetic data provide strong evidence that H3 K27M impairs, but does not abolish, PRC2 activity. Enhancers RNAi lines, particularly those targeting core Polycomb components such as E(z), Su(z)12, and members of cPRC1, vPRC1, PRC2 and PhoRC, exacerbated the phenotypes of H3 K27M-expressing tissues. These findings are consistent with prior observations in both *Drosophila* and mammalian systems, in which PRC2 activity is partially retained in the presence of H3 K27M. The exacerbation of phenotypes upon enhancer knockdown suggests that residual PRC2 or PRC1 activity is functionally important and may play a buffering role that prevents further transcriptional dysregulation. These findings mirror data from human DMG and PFA ependymoma, in which complete loss of Polycomb activity is incompatible with tumor cell viability (Bender et al. 2013; Mohammad et al. 2017; Piunti et al. 2017; Harutyunyan et al. 2019; Jain, Rashoff, et al. 2020; Sarthy et al. 2020; Chaouch et al. 2021). Thus, residual Polycomb activity is both biologically relevant and potentially therapeutically targetable.

### Suppressors identify transcriptional networks that cooperate with PRC2 inhibition to drive pathology

In contrast to enhancers, suppressors identify pathways that are required for the detrimental effects of H3 K27M. Remarkably, suppression occurred without restoration of H3K27me3 levels, demonstrating that impaired PRC2 activity alone is insufficient to produce developmental arrest. Many of the strongest suppressors we identified deposit marks of active chromatin: Trx (H3K4me1/me2), Ash1 (H3K36me2), and Set2 (H3K36me3). These findings are consistent with longstanding evidence that H3K4 and H3K36 methylation antagonize Polycomb-mediated repression (Zhang et al. 2015; Hyun et al. 2017). Indeed, Trithorax group proteins were first identified as suppressors of dominant Polycomb group mutants in *Drosophila* (Kassis et al. 2017), and Ash1 was identified as a suppressor of H3 K27M expression in the *Drosophila* eye (Chaouch et al. 2021). Biochemical and genetic data have suggested that methylation of H3K4 and H3K36 antagonize Polycomb group activity (Schmitges et al. 2011; Lewis et al. 2013; Jain, Khazaei, et al. 2020). Although previous studies had only identified Ash1 as a suppressor of H3 K27M in the *Drosophila* eye (Chaouch et al. 2021), our screen also identified Set2 and two additional H3K36 methyltransferases (NSD and CG4565), underscoring the central role of H3K36 methylation in modulating H3 K27M-driven phenotypes. These effects were not mediated through global restoration of PRC2 activity but rather through modulation of the transcriptional consequences of its loss. We demonstrate that reducing H3K36 methylation does not restore global H3K27me3 levels and that Ash1 depletion lowers the expression levels of genes upregulated by H3 K27M expression (Figure 5B and data not shown). Together with prior findings, these results support a model in which a balance between H3K36 methylation and H3K27me3 is essential for maintaining gene-expression levels required for normal development.

Along with previously identified Trithorax group proteins, we identified Asx as a strong suppressor that restores normal development to flies expressing H3 K27M. RNA-seq in wing discs depleted of A*sx, Ash1,* and *Trx* identified highly overlapping transcriptional targets, suggesting shared mechanisms of action. Notably, Asx is a subunit of the PR-DUB complex that removes PRC1-mediated H2AK118ub (H2AK119ub in humans) (Scheuermann et al. 2010). Although we did not identify Calypso (BAP1 in mammals), likely due to developmental toxicity from its knockdown, our data suggest that Asx plays a transcription-promoting role in the wing. Nonetheless, we cannot rule out that the function identified is independent of the deubiquitination of H2A. The relationship between PR-DUB-mediated deubiquitination and Polycomb-mediated repression remains unresolved, with conflicting evidence for roles in both repression and activation (Balasubramani et al. 2015; LaFave et al. 2015; Conway et al. 2021). Our findings support a model in which, in some developmental contexts, PR-DUB components, such as Asx, promote gene expression in a manner that is functionally antagonistic to PRC2.

### Nup153 suppression reveals an additional chromatin regulatory axis distinct from Trithorax group proteins

Amongst the strong suppressors identified was Nup153, a dynamic nucleoporin known to bind to euchromatin and mediate both gene activation and repression (Vaquerizas et al. 2010; Jacinto et al. 2015; Ibarra et al. 2016; Toda et al. 2017; Kadota et al. 2020; Pascual-Garcia and Capelson 2021; Capelson 2023). In mammalian tissue culture, Nup153 has been implicated in PRC1-mediated silencing (Jacinto et al. 2015). In flies, it has been suggested to act as a transcriptional activator, but its molecular functions remain largely enigmatic (Vaquerizas et al. 2010). Our identification of Nup153 as a strong suppressor suggests it might function in promoting gene expression, similar to other identified suppressors. Although RNAi of Nup153 RNAi strongly suppressed the phenotypes caused by expression of H3 K27M and EZHIP, it produced largely distinct transcriptional consequences from those of Trx, Ash1, and Asx. However, Nup153 itself was among the small set of 31 genes downregulated by all four suppressors, raising the possibility that downregulation of Nup153 may be a common feature required for suppression. Given its context-dependent chromatin functions, Nup153 may facilitate or stabilize gene-expression programs that become deleterious specifically in the context of PRC2 inhibition. Its downregulation in multiple suppressor backgrounds suggests that H3 K27M-expressing cells may require wild-type levels of Nup153 to sustain the transcriptional rewiring associated with PRC2 inhibition. This potential dependency raises the intriguing possibility that Nup153 represents a novel therapeutic vulnerability in PRC2-inhibited tumors.

### A model for PRC2 inhibition: oncoprotein phenotypes arise from regulatory imbalance

Our findings support a model in which PRC2 inhibition by H3 K27M disrupts a critical balance between chromatin repression and transcriptional activation, leading to developmental arrest. This disruption arises not solely from the loss of H3K27me3 but from the unopposed activity of transcriptional activators that promote gene expression at previously silenced loci (Figure 6). In this framework, PRC2 inhibition by H3 K27M poises chromatin for expression by eliminating repressive spreading of H3K27me3, but gene activation requires additional inputs from proteins such as Trx, Ash1, and Asx. Knockdown of these factors restores the critical gene-regulatory balance, allowing normal development to proceed. Such a model would explain why PRC2 inhibition is necessary, but not sufficient, to drive H3 K27M phenotypes, and why suppressors can reverse aberrant development even in the absence of restored H3K27me3.

The regulatory imbalance we uncovered is not unique to H3 K27M as there is evidence that gene-regulatory imbalances drive other forms of cancer (Biegel et al. 1999; Wang et al. 2018; Wang et al. 2019; Drosos et al. 2022). Many cancers are driven by mutations in transcriptional activators, including suppressors that emerged in our screen (Corral et al. 1996; Faber et al. 2009; Mills 2010; Zhu et al. 2016; Rogawski et al. 2021). In these contexts, the repressive function of PRC2 may become essential to counteract hyperactivated gene expression programs. Thus, our work not only elucidates the mechanisms by which H3 K27M and EZHIP disrupt development but also reveals broader principles of chromatin regulation that are relevant to cancer biology. Together, our genetic screen has uncovered previously unappreciated roles for conserved chromatin regulators in mediating the phenotypes caused by inhibition of allosterically activated PRC2, with important implications for our understanding of the pediatric cancers caused by H3 K27M and EZHIP expression. These findings underscore the importance of studying chromatin dynamics *in vivo* and suggest that therapeutic strategies targeting transcriptional activators or nucleoporins like Nup153 may be effective in mitigating the consequences of PRC2 inhibition in pediatric gliomas. Future studies defining the chromatin landscapes and protein interaction partners responsible for suppression will provide deeper mechanistic understanding and potentially inform new avenues for intervention.

## Materials and Methods

### Fly strains/husbandry

All stocks were grown on molasses food at 25°C unless otherwise noted. N-terminally FLAG-tagged *EZHIP*, *EZHIP M406E, H3.3 K27M* or *H3.3 K27R* were cloned into pUASt-attB (DGRC #1419) using PCR, restriction digest and ligation. Transgenes were integrated into the *M{3xP3-RFP.attP}ZH-86Fb* locus on chromosome three (Bloomington *Drosophila* Stock Center (BDSC) #24749), or into the *PBac{yellow[+]-attP-9A}VK14* locus on chromosome two (BDSC #9733) using PhiC31 integrase-mediated recombination with fluorescence marker removed (Best Gene, Chino Hills, CA).

### Selection of lines for RNAi screen

FlyMine online software was used to query all *Drosophila* genes with chromatin-related functional annotations.(Lyne et al. 2007) Among these, ten chromatin-related gene ontology (GO) terms were selected for further analysis (Supplemental Figure 2A). siRNA and miRNA-related genes were removed given the RNAi-based screening approach used. Genes were excluded if not expressed in a wing imaginal disc-derived cell line based on available RNA-seq data ML-DmD21 (DGRC Stock #86) (Stoiber et al. 2016). Conservation of genes between *Drosophila* and humans was queried with *Drosophila* Integrative Ortholog Prediction Tool (DIOPT), an integrated tool that uses nine ortholog predictors (Hu et al. 2011). Only genes deemed to be highly conserved according to DIOPT were included in our screen.

All RNAi lines included in our screen were generated by the Transgenic RNAi Project (TRiP) from the Harvard *Drosophila* RNAi Screening Center (DRSC) (Perkins et al. 2015). TRiP stocks were generated over multiple generations and used different cloning strategies to generate dsRNA transgenes for each target gene. Vectors vary by dsRNA expression level, and production of long- or short-hairpin dsRNA, which have weaker or stronger average target gene knockdown, respectively. We considered TRiP stocks “strong” or “weak” depending on their expressed dsRNA hairpin length. We used the Updated Targets of RNAi Reagents (UP-TORR) tool to identify available TRiP stocks (Hu et al. 2013). Depending on reagent availability, we ordered long- and short-hairpin RNAi lines for every gene in our screen. Reagent availability was determined using the DRSC/TRiP Functional Genomics Resources Lookup (Hu et al. 2013). Only somatically expressing RNAi vectors were included in our screen (dsRNA constructs cloned into VALIUM1, VALIUM10, or VALIUM20).

### Immunostaining

For imaginal wing disc immunostaining, flies carrying *UASt-EZHIP*, *UASt-H3 K27M*, *UASt-H3 K27R*, or Harvard Transgenic RNAi project (TRiP) lines were crossed to *nub*-*GAL4* (II); *UAS*-*Dcr-2 (X)* (BDSC#25754) for wing-specific transgene expression. Wing imaginal discs were harvested from crawling third-instar larvae for immunostaining. For eye-antennal disc immunostaining and enhancer/suppressor screening in the eye, a recombinant *ey*-*GAL4*,*GMR*-*GAL4 (II)* line was used to drive transgene expression (gift from the Lab of Dr. Nansi-Jo Colley). H3 K27M was co-expressed with RNAi lines to perform enhancer/suppressor eye screen.

Wing and eye-antennal imaginal discs were dissected from crawling third-instar larvae into ice-cold 1X PBS. Discs were fixed for 30 minutes at room temperature in a 4% formaldehyde-1X PBS solution, then permeabilized with 1X PBS + 0.1% Triton X-100 (PBST). After permeabilization, discs were blocked with PBST + 1% BSA (PAT) for ten minutes at room temperature and incubated in PAT overnight at 4°C with the following primary antibodies: rabbit anti-H3K27me3 (1:1600) (Cell Signaling Technology #9733S), and mouse M2 anti-FLAG (Sigma #F1804). The following day, discs were washed in PBST and blocked with PBST + 2% normal goat serum for ten minutes prior to addition of secondary antibodies. Discs were incubated with secondary antibodies (1:2000) in PBST + 2% normal goat serum for four hours at room temperature. For FLAG staining, goat anti-mouse 488 DyLight conjugated secondary antibody was used (Fisher Scientific #35502) was used. For H3K27me3, goat anti-rabbit conjugated Alexa Fluor 594 (Fisher Scientific #A-11001) secondary antibody was used. DAPI (ThermoFisher #D1306) was added to secondary antibody solution (1:2000) for 5 minutes before final washes and mounting. Discs were imaged at 10X using a Nikon Ti2-E epifluorescent microscope. Final images were processed using imageJ v1.52.

### Adult wing and eye imaging

Adult flies were imaged while anesthetized. For wing imaging, flies were placed in 70% ethanol at −20°C for at least 15 minutes, but up to six weeks. Once removed from ethanol, wings were dissected from flies in 1x PBS solution and mounted in 70% glycerol. All images were acquired with an OMAX 18MP USB 3.0 C-Mount camera placed in the eyepiece of a dissecting microscope at 4X magnification. Camera operated with ToupLite imaging software on laptops. The following RNAi lines were used to generate adult fly wing images in this manuscript: *E(z)* RNAi (BDSC #31617), *trx* RNAi (BDSC #31092), *CycC* RNAi (BDSC #33753), and *Usp7* RNAi (BDSC #34708). The following RNAi lines were used to generate adult eye images: *E(z)* RNAi (BDSC #31617), *trx* RNAi (BDSC #31092), and *ash1* RNAi (BDSC #33705).

### Enhancer/suppressor screen

Fly lines were generated for enhancer/suppressor screen using recombination of *nub*-*GAL4 (II)*; *UAS*-*Dcr-2 (X)* (BDSC #25754) with *UASt-EZHIP*, *UASt-H3 K27M*, and *UASt-H3 K27R*. Recombination was confirmed using PCR screening, wing phenotyping, and immunostaining. Every RNAi line was crossed to the following fly lines to generate RNAi and screen scores: *nub*-*GAL4 (II)*; *UAS*-*Dcr-2 (X),* and the recombinant *nub*-*GAL4, UASt-H3 K27M(II)/CyO*; *UAS*-*Dcr-2 (X)*, respectively. RNAi and screen scores were generated by two independent researchers who were blinded to the identity of the RNAi target genes. Only RNAi lines classified as enhancers by both researchers were included in the final analysis. Every enhancer and suppressor was crossed to *nub*-*GAL4, UASt-H3 K27R/CyO (II)*; *UAS*-*Dcr-2 (X)* to control for nonspecific effects of histone transgene overexpression. All suppressors were additionally crossed to *nub*-*GAL4, UASt-EZHIP/CyO (II)*; *UAS*-*Dcr-2 (X)*. All recombinant lines were viable and fertile.

### RNA-seq and CUT&RUN transgene expression

We generated a fly line to ubiquitously express H3 K27M or H3 K27R, and TRiP RNAi lines in a temperature-sensitive manner. Briefly, we used recombination of *Act5C-GAL4 (II)* (BDSC #25374) and *alphaTub84B-GAL80ts (II)* (BDSC #7019). Recombination was confirmed using PCR and immunostaining. *Act5C-GAL4, alphaTub84B-GAL80ts (II)* was crossed to fly lines carrying *UASt-H3 K27M* or *UASt-H3 K27R*. Animals were reared at 25°C, a temperature at which GAL80 represses GAL4. Plates with molasses and yeast paste were exchanged at three-hour intervals to stage discs at time of dissection. Embryos were aged on plates for 24 hours and picked into vials as first-instar larvae. Larvae were reared until 44 hours after egg laying (AEL), then shifted to 32°C to inactivate GAL80 and express transgenes for 72 hours.

### RNA-sequencing

Imaginal wing and eye-antennal discs were harvested from crawling third-instar larvae. Larvae were staged to within 116 and 119 hours after egg laying. For each replicate, ten discs were dissected into ice cold 1X PBS solution. Three biological replicates were dissected for each genotype. After dissection, discs were incubated in Trizol (Invitrogen #15596026) for 5 minutes to dissolve tissue, then frozen at −20°C. RNA was isolated using standard Trizol RNA isolation procedure, and libraries were prepared using the TruSeq RNA sample prep kit v2 (Ilumina RS-122-2002). 75-base-pair reads were obtained using an Illumina NextSeq500 High-Throughput Sequencer. Sequencing was performed at the Northwestern University Sequencing (NUSeq) Core Facility. Suppressor RNAi lines used for RNA-seq included: *trx* RNAi (BDSC #31092), *ash1* RNAi (BDSC #33705), *Asx* RNAi (BDSC #31192), and *Nup153* RNAi (BDSC #32837).

Discs harvested for RNA-seq had the following genotypes:

– *Act5C-GAL4, alphaTub84B-GAL80ts/UASt-H3 K27M*
– *Act5C*-*GAL4, alphaTub84B-GAL80ts/UAS-tH3 K27M; Asx/trx/ash1/Nup153 RNAi/*
– *Act5C*-*GAL4, alphaTub84B-GAL80ts/UASt-H3 K27R*
– *Act5C*-*GAL4, alphaTub84B-GAL80ts/UASt-H3 K27R; Asx/trx/ash1/Nup153 RNAi/+*

### RNA-seq analysis

RNA-seq data was aligned to the dm6 *Drosophila melanogaster* genome using HISAT v2.1.0 (Kim et al. 2015). Multi-mapping reads were excluded from further analysis. featureCounts v1.5.3 was used to generate a table with reads assigned to annotated dm6 genes using UCSC annotation r6.45 (Liao et al. 2014). Read counts were used to determine differentially expressed genes using DESeq2 v1.14.1 (Love et al. 2014). Genes with an adjusted p-value <0.05 and log_2_ fold change >1.5 were considered differentially expressed. Genes with <50 reads across all samples were excluded from analysis. Differential expression for selected groups of dysregulated genes determined based on log_2_RPKM-normalized reads. Volcano plots and violin plots generated in RStudio v1.4.1106 using ggplot2 v3.4.2. Heatmaps generated using pheatmap v1.0.12. Venn Diagrams generated using DeepVenn (arXiv:2210.04597). Mean-average (MA) plots generated using ggpubr v0.5.0 or ggplot2 v3.4.2.

### Cleavage under targets, release under nuclease (CUT&RUN)

Wing imaginal discs were harvested from crawling third-instar larvae between 116 and 119 hours AEL. Two biological replicates were used to assay H3K27me3 for each genotype, and one replicate was used in IgG control. For each replicate, twenty discs were dissected into ice-cold PBS. Approximately 50,000 cells are in a third-instar larval wing disc, amounting to one million cells for each replicate. Intact wing discs were used for the entire protocol. Samples were washed three times in Wash Buffer (20mM HEPES pH 7.5, 150mM NaCl, 0.5 mM spermidine (Sigma S0266-1G), and Pierce EDTA-free Protease Inhibitor (ThermoFisher PIA32955)) and incubated with activated concanavalin-A (ConA) paramagnetic beads (EpiCypher SKU: 21-1401) for ten minutes in PCR strip tubes. Samples were resuspended in cold Antibody Buffer (Wash Buffer + 0.05% digitonin (Sigma Millipore 300410250MG) and 2mM EDTA (Fisher S311500)). 2 uL of SNAP-CUTANA K-MetStat nucleosomes (EpiCypher SKU: 19-1002) diluted 1:100 in Wash Buffer were added to each sample for spike-in normalization. Antibody was added to samples at a 1:50 concentration and incubated overnight at 4°C (Rabbit anti-H3K27me3 (Cell Signaling Technology #9733S) or rabbit IgG control (ThermoFisher #10500C)). The following day, discs were washed with Digitonin Buffer (Wash Buffer + 0.05% digitonin (Sigma Millipore 300410250MG)), then incubated with pAG-MNase (EpiCypher SKU: 15-1016) for ten minutes to allow protein A/G binding to antibody. 1 uL of cold 100 mM CaCl_2_ was added to samples to activate pAG-MNase and incubated on nutator for 2 hours at 4°C. Reaction was quenched with Stop Buffer (340 mM NaCl, 20 mM EDTA, 4 mM EGTA, 50 µg/mL RNase A, 50 µg/mL Glycogen). Chromatin was eluted from samples at 37°C for ten minutes, then DNA was purified with the Qiagen MinElute Reaction Cleanup Kit (Qiagen #28204). Libraries were prepared using the NEBNext Ultra II DNA Library Prep Kit for Illumina (NEB E7645L) and sequenced on the Illumina NovaSeq6000 High-Throughput Sequencer at the University of Wisconsin Biotechnology Center (UWBC). Sequencing produced 150-base-pair paired-end reads.

Discs harvested for CUT&RUN had the following genotypes:

– *Act5C-GAL4, alphaTub84B-GAL80ts/UASt-H3 K27M*
– *Act5C*-*GAL4, alphaTub84B-GAL80ts/UASt-H3 K27R*

### CUT&RUN & ChIP analysis

Read quality was assessed using FASTQC (v0.11.9).(Babraham Bioinformatics - FastQC A Quality Control tool for High Throughput Sequence Data) Adapters and low-quality bases were removed using Trimmomatic (v0.39.29) (Bolger et al. 2014a). Reads were mapped to the dm6 genome assembly (dos Santos et al. 2015) using Bowtie2 (Langmead and Salzberg 2012). Unmapped, multiply aligning, mitochondrial, and scaffold reads were removed. Nonspecific CUT&RUN signal was subtracted from H3K27me3 samples using IgG control for each genotype. MACS2 was used to identify broad H3K27me3 peaks after merging replicates for each genotype (Zhang et al. 2008). Standard MACS2 parameters were used to call narrow peaks for published Ph ChIP data (Loubière et al. 2016). Antibody specificity was assessed using percentage of on-target spike-in reads compared to total spike-in reads. Merged alignment files were normalized using a combination of RPKM and spike-in. Spike-in normalization based on percentage of reads mapping to all barcoded nucleosomes. Integrated Genomics Viewer (v2.12.3) was used for visualization of normalized bigWig files (Thorvaldsdóttir et al. 2013). ΔH3K27me3 bigWig files were generated using bamCompare function in deepTools (v3.4.1). RPKM, spike-in normalized read counts for H3 K27R samples were subtracted from normalized H3 K27M read counts. Heatmaps were generated using deepTools (v3.4.1) with RPKM, spike-in normalized bigWig files (Ramírez et al. 2016). Analysis of H3K27me3 enrichment at wing disc based on differentially expressed gene sets using rtracklayer (v3.16). Genomic annotations were derived from UCSC annotated dm6 genome r6.45.

## Acknowledgments

We acknowledge the Harvard Transgenic RNAi Project and the Bloomington *Drosophila* Stock Center for generating RNAi lines and providing fly lines. We also acknowledge the University of Wisconsin-Madison Biotechnology Center and the NUSeq Core Facility for sequencing. Experiments were supported by grants from the National Institutes of Health: F30CA260987 (SDK), R01CA266861 (PWL), P01CA196539 (PWL), and the UW Comprehensive Cancer Center Support Grant (MMH and PWL).

## Supplemental Tables

**Supplemental Table 1:** RPKM normalized RNA-seq data. Column 1 is the gene identifier. Column 2 is the gene symbol. Each subsequent column is labelled with the individual datasets

**Supplemental Table 2:** Genes with increased expression upon H3 K27M expression as compared to H3 K27R in the wing (Tab 1) or eye (Tab 2) imaginal discs.

**Supplemental Table 3:** Results of the RNAi screen in combination with H3 K27M (Tab 1), H3 K27R (Tab 2) or EZHIP (Tab 3) expression.

**Supplemental Table 4:** Target genes whose RNAi knockdown modified the H3 K27M phenotype. Associated strains from the Bloomington Drosophila Stock Center and human orthologs are listed. Green denotes suppressors and blue denotes enhancers (as indicated).

## Notes

### Competing Interest Statement

The authors have declared no competing interest.

