## Supplemental Figures for "Investigating oncoprotein-mediated chromatin dysregulation in *Drosophila melanogaster* uncovers novel modifiers of the developmental impact of H3 K27M and EZHIP"

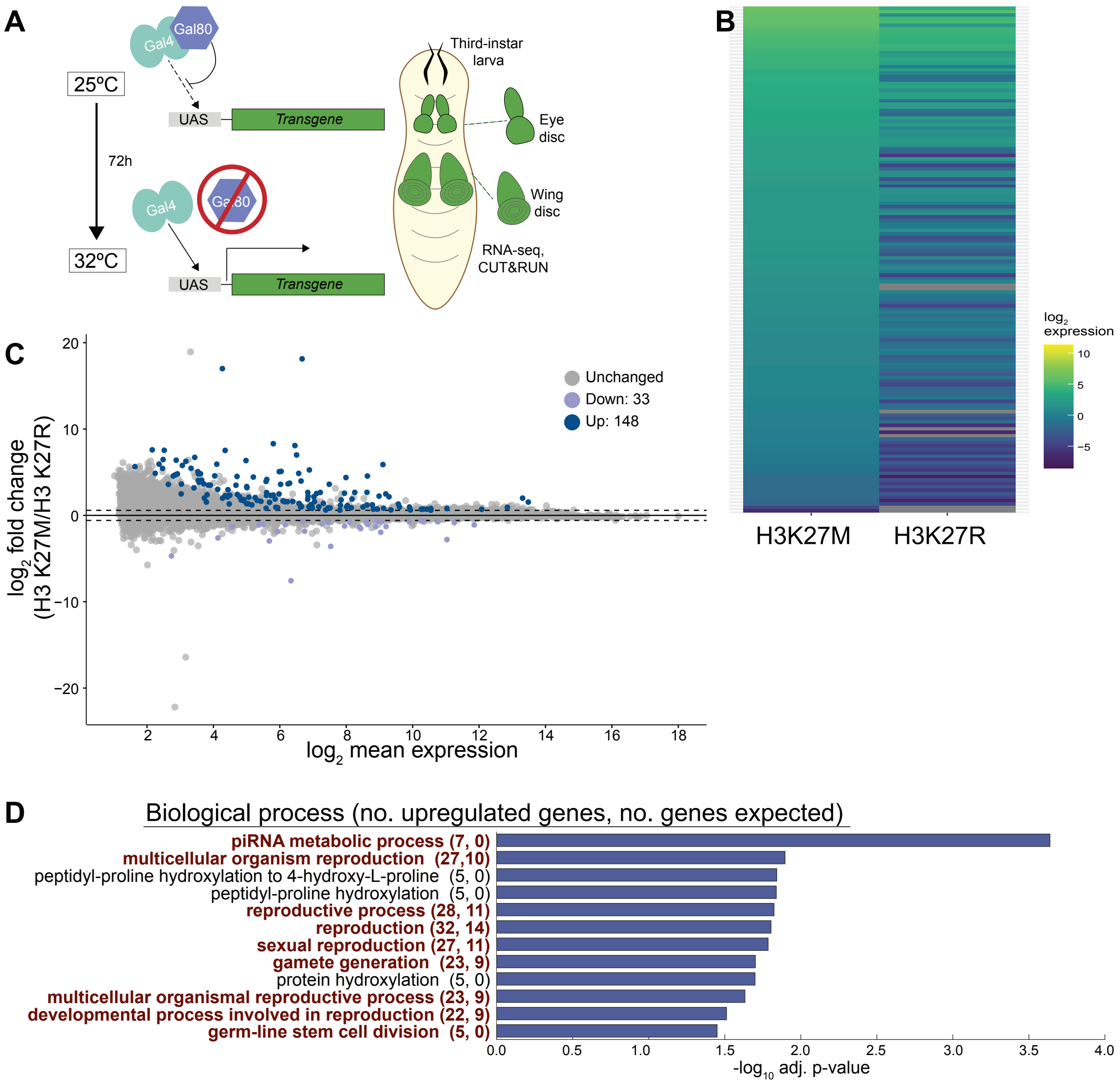


**Supplemental Figure 1. H3 K27M promotes expression of germline-related genes in the wing disc.** A. Schematic of transgene expression system used for RNA-seq and CUT&RUN. Transgenes expression under control of ubiquitously expressed *actin-Gal4* driver. *tubulin-Gal80* is a ubiquitously expressesd, temperature-sensitive Gal4 inhibitor. At 25ºC, Gal80 is active and represses Gal4-mediated transgene expression. At 32ºC, Gal80 is inactivated, permitting transgene expression. 72 hours after shifting developing animals to 32ºC, wing imaginal discs were harvested from third-instar larvae and used in RNA-seq and CUT&RUN. B. Expression ($log_2$ RPKM) of the 148 genes upregulated in wing discs expressing H3 K27M compared to H3 K27R (adjusted p-value < 0.05; fold change > 1.5). C. MA plot highlighting differentially-expressed genes in H3 K27M wing discs compared to H3 K27R. Upregulated genes are denoted by blue dots. Upregulated genes are shown in dark blue. Downregulated genes are in lavender. (adjusted p-value < 0.05, fold change > 1.5). Gray dots designate genes whose expression was unchanged. D. Gene ontology (GO) analysis for genes upregulated by H3 K27M in wing discs. Number of genes upregulated by H3 K27M listed first in parentheticals, followed by expected number of upregulated genes . $-log_{10}$ adj. p-value plotted. Genes involved in germline related processes are highlighted in magenta and bolded.

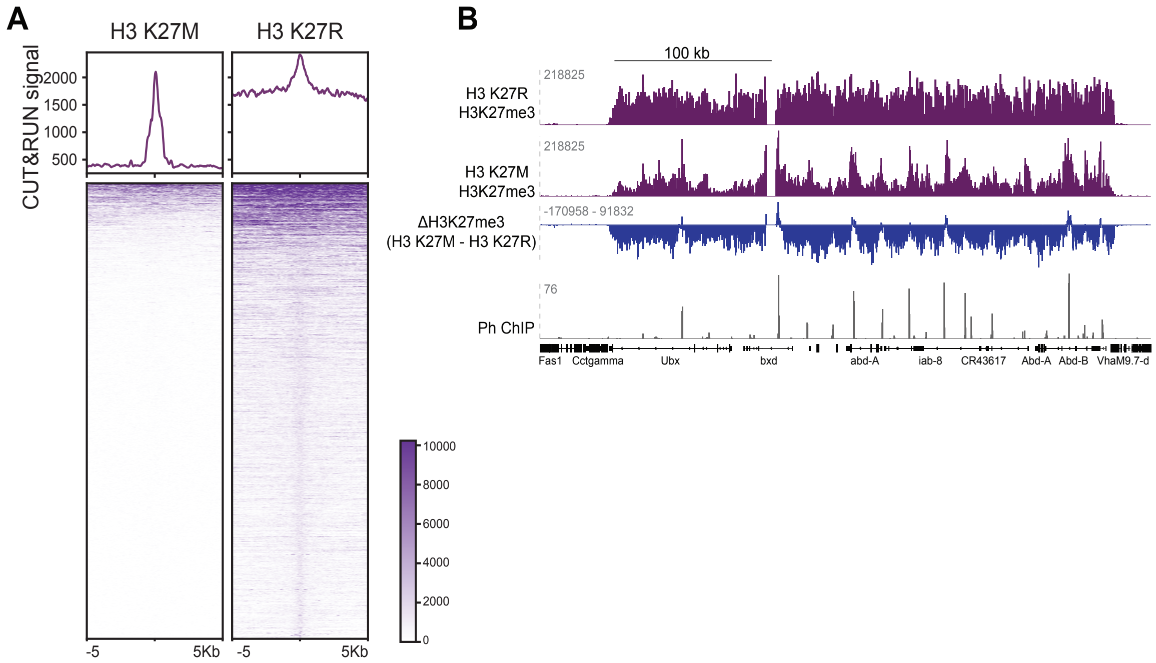


**Supplemental Figure 2. H3 K27M expression inhibits spreading of H3K27me3.** A. Cleavage under targets, release under nuclease (CUT&RUN) assayed H3K27me3 in wing discs expressing H3 K27M or H3 K27R. Top: Metaplots of H3K27me3 enrichment (z-score and spike-in normalized) at all H3K27me3 peaks in wing discs expressing H3 K27R. Bottom: Heatmaps of normalized H3K27me3 enrichment at peaks overlapping H3K27me3 peaks in H3K27R wing discs, sorted by highest to lowest signal intensity and centered on peak in H3 K27R-expressing discs. B. Top, purple: Genome browser tracks showing normalized H3K27me3 enrichment at *Ubx* and *Abd-A/Abd-B*, well-known targets of Polycomb mediated repression. Middle, blue: difference in normalized H3K27me3 enrihcment between wing discs expressing *H3 K27M* and *H3 K27R*. Regions with positive signal indicate increased H3K27me3 in H3 K27M wing discs. Bottom, gray: Ph enrichment in wing discs, based on published ChIP-seq (Loubiere et al. 2016) Regions with positive signal in ΔH3K27me3 tend to overlap Ph peaks.

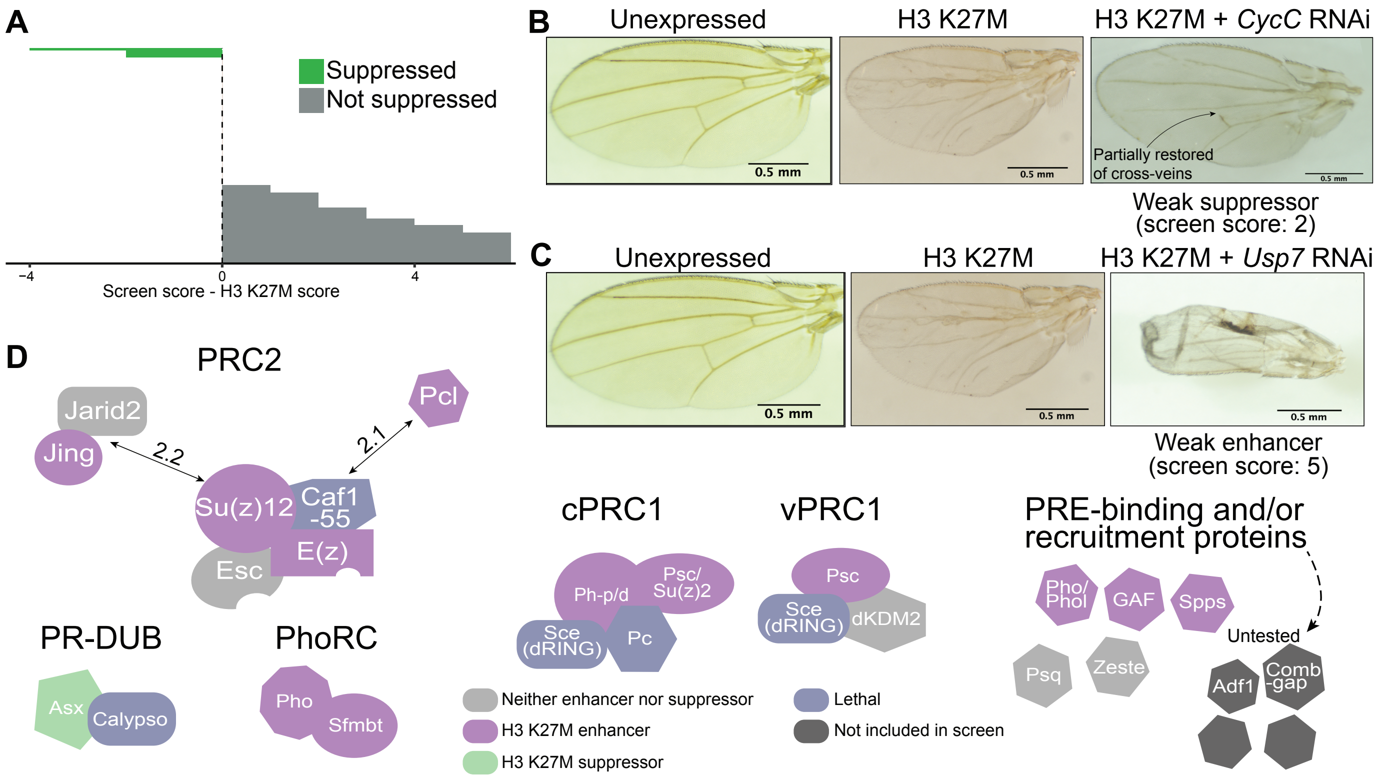


**Supplemental Figure 3. Knockdown of conserved chromatin modifiers revealed novel enhancers and suppressors of H3 K27M phenotypes.** A. Difference between screen score and H3 K27M-alone score for every RNAi line in screen. If screen score minus the H3 K27M score produces a negative value, the RNAi line was classified as a suppressor. -4 indicates strong suppression, as H3 K27M phenotype was assigned 4 points and wild-type wings were assigned 0. B. Example of a weak suppressor. Co-expression of H3 K27M with *CycC* RNAi eliminated the wrinkled phenotype found in H3 K27M wings but only partially restored normal wing veins (2 points). CycC was thereby assigned a screen score of 2. C. Example of a weak enhancer. Co-expression of H3 K27M with *Usp7* RNAi produced wings with wrinkling (2 points) and recognizable, albeit mispatterned wing veins (2 points) like H3 K27M. However, there was a mild to moderate size decrease compared to H3 K27M (score of 1) for a total score of 5. *Usp7* RNAi produced an RNAi score of 0. D. Summary of Polycomb proteins included in screen. Polycomb proteins were the most enriched functional group among enhancers. A number of the Polycomb proteins not called as enhancers were lethal (in blue) and therefore wing phenotypes were not able to be scored.

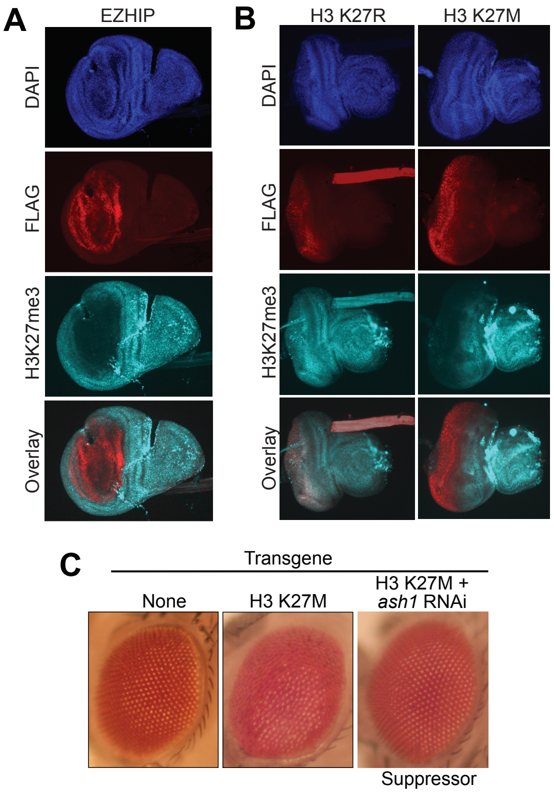


**Supplemental Figure 4. H3 K27M and EZHIP phenotypes are suppressed in multiple tissues independent of PRC2 inhibition.** A. Immunostaining for FLAG (transgene) and H3K27me3 in wing imaginal discs dissected from third-instar larvae. EZHIP was expressed under control of *nubbin-Gal4*. DAPI stains nuclei. B. Immunostaining for FLAG (transgene) and H3K27me3 in eye-antennal imaginal discs dissected from third-instar larvae. Transgenes expressed under control of *eyeless,GMR-Gal4* driver. C. H3 K27M disorganizes photoreceptors on surface of eye. Co-expression of H3 K27M with *ash1* RNAi rescues wild-type eye development.

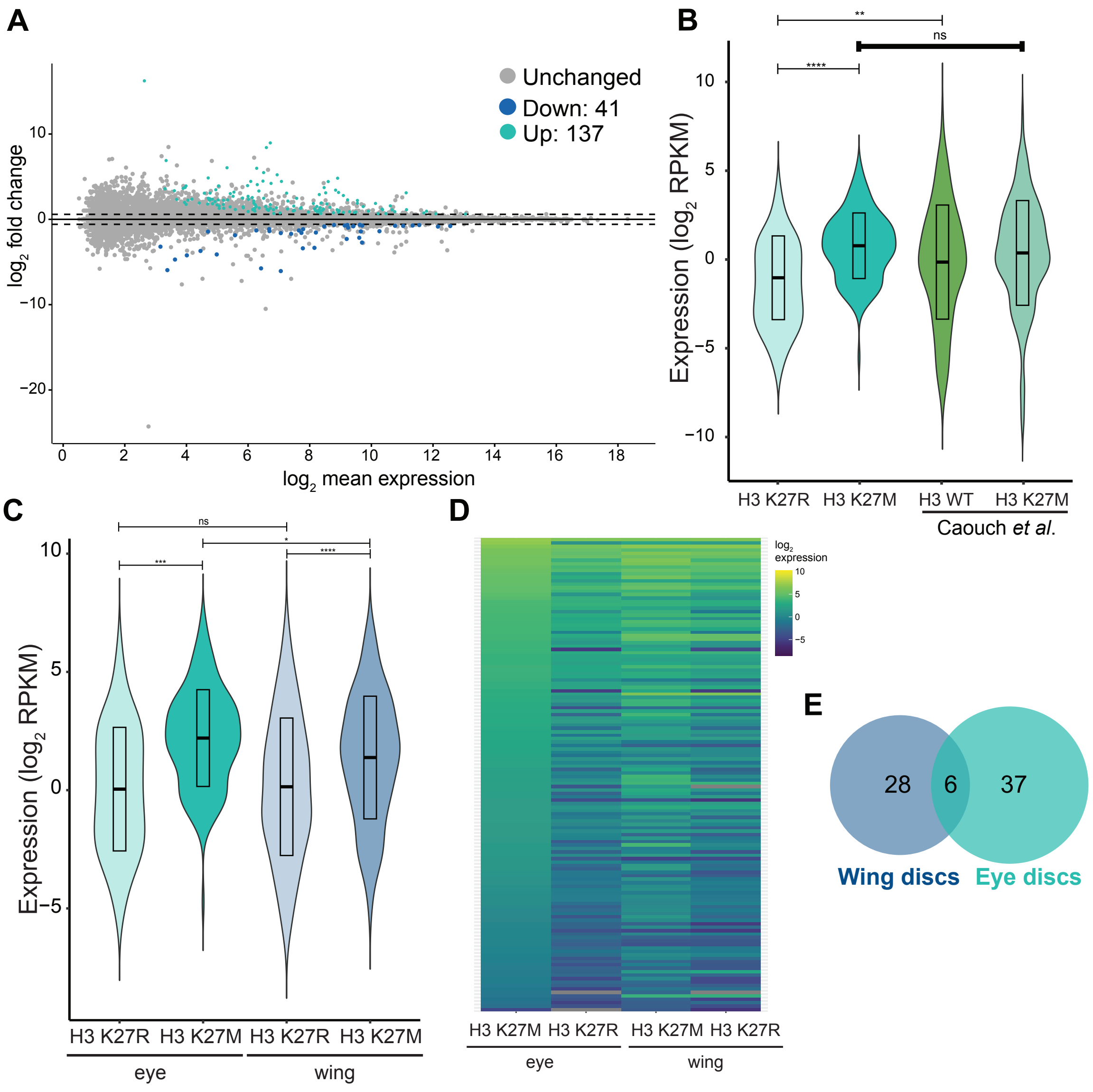


**Supplemental Figure 5. H3 K27M induces similar transcriptional changes in the wing and eye discs.** A. MA plot of genes in eye-antennal discs expressing H3K27M as compare to H3 K27R controls. Light blue dots designate upregulated or downregulated genes (adjusted p-value < 0.05, fold change > 1.5). Dark blue dots are downregulated genes. Gray dots designate genes with non-significant changes. B. Violin plot showing average expression (RPKM, $log_2$) of the 137genes upregulated by H3 K27M in eye-antennal discs compared to H3 K27R. Expression is shown for genotypes and tissues indicated below. For comparison, data from Cauch *et al*. is shown. Differences are largely due to differences in H3 WT controls versus H3 K27R controls in the two different data sets. ns = not significant (adjusted p-value > 0.05). ****adjusted p-value < 0.0001. (one-way ANOVA) C. Violin plot showing average expression (RPKM, $log_2$) of the 137 genes upregulated by H3 K27M in eye-antennal discs compared to H3 K27R. Expression of the same gene set in the wing discs are also shown (as indicated below). ns = not significant (adjusted p-value > 0.05). ****adjusted p-value < 0.0001. (one-way ANOVA) I. Heat map of the $log_2$ expression of 137 genes upregulated by H3 K27M eye-antennal discs as compared to H3 K27R discs. Expression of the same genes in the wing disc is also shown (as indicated below). Genes ordered by average expression in H3 K27M eye-antennal discs.C. Venn diagram of the overlap among genes downregulated by H3 K27M in wing and eye discs.

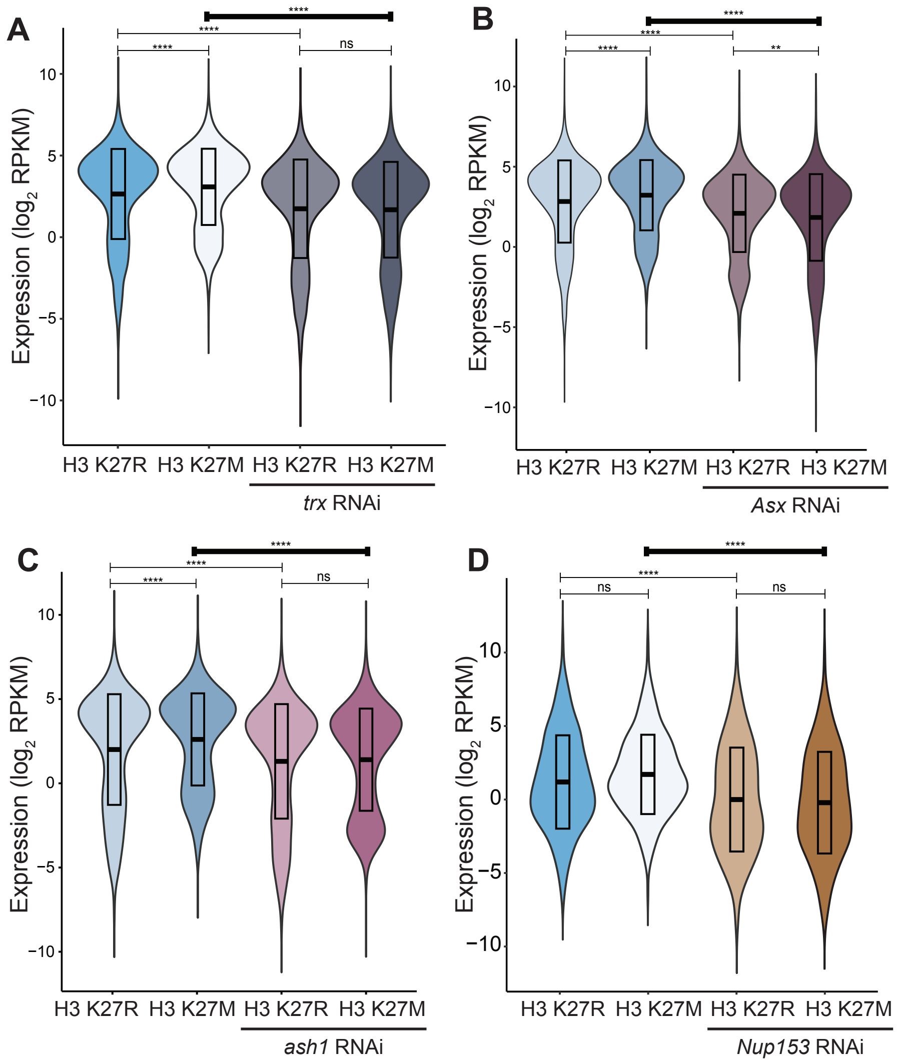


**Supplemental Figure 6. RNAi of H3 K27M suppressors decreases transcript levels of genes whose levels increased upon H3 K27M expression in wing discs.** A-D. Violin plots showing average expression ($log_2$ of RPKM) of genes downregulated in wing discs expressing H3 K27M and the suppressor RNAi indicated as compared to H3 K27M discs alone. ns = not significant (adjusted p-value > 0.05). ****adjusted p-value < 0.0001 (one-way ANOVA).

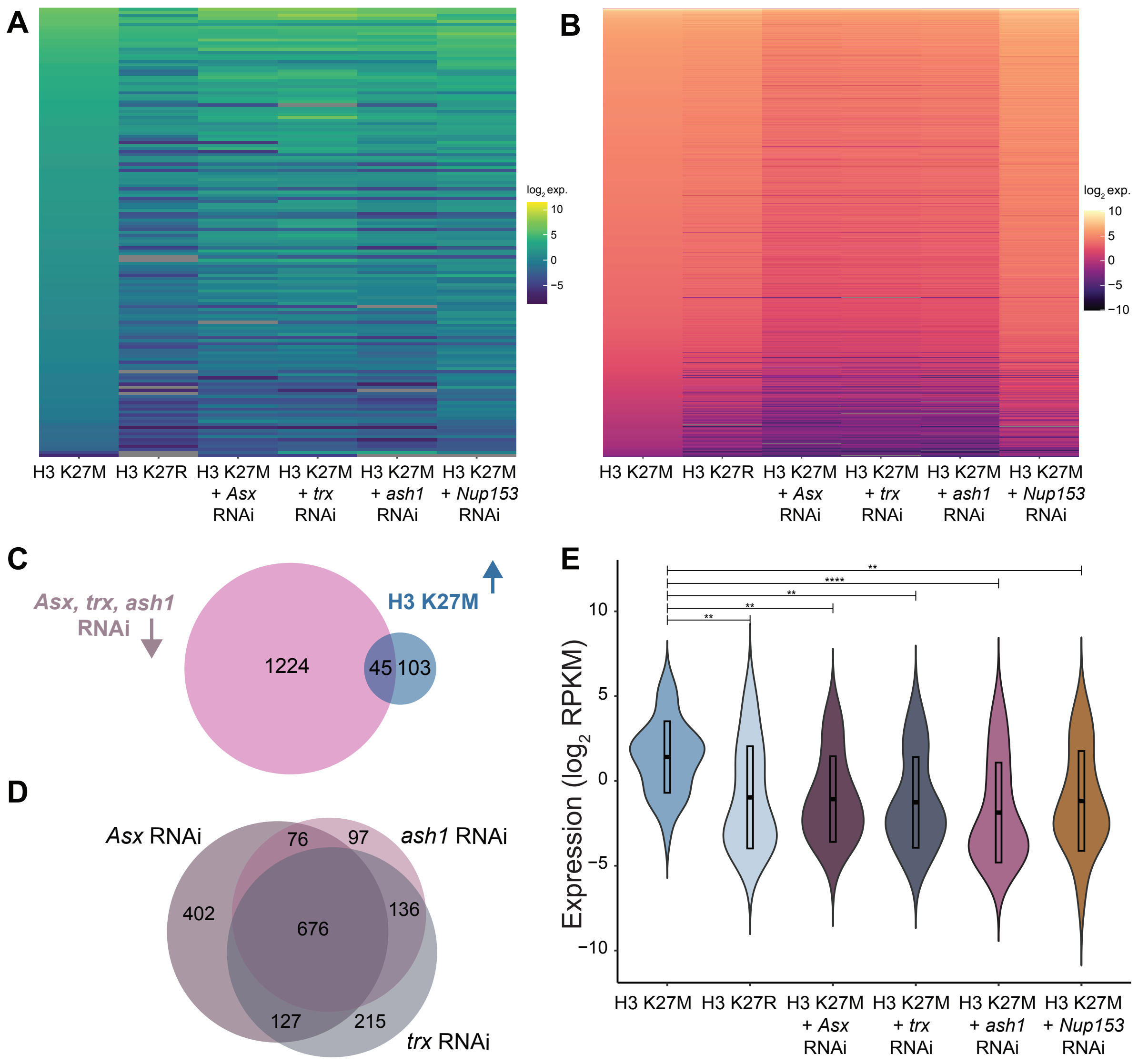


**Supplemental Figure 7. Expression of a subset of genes is restored to wild-type levels in suppressors.** A. Heat map of RNA expression levels (log2 of RPKM) for the 148 genes upregulated by H3 K27M in wing discs compared to H3 K27R for the genotypes indicated below. B. Heat map of RNA expression levels ($log_2$ RPKM) for the 1269 genes downregulated by *Asx*, *trx*, and *ash1* RNAi in H3 K27M-expressing wing discs. C. Venn diagram showing overlap between the 148 genes upregulated by H3 K27M as compared to H3 K27R and the 1269 genes downregulated *Asx*, *trx*, and *ash1* RNAi in H3 K27M-expressing wing discs. D. Venn diagram showing overlaps between genes downregulated by *Asx*, *trx*, and *ash1* RNAi when co-expressed with H3 K27R compared to H3 K27R alone. E. Violin plot of the average expression ($log_2$RPKM) of the 31 genes downregulated upon RNAi of all four suppressors. For comparison, expression levels are also shown for H3 K27M and H3 K27R-expressing wing discs alone. ns not significant (adjusted p-value > 0.05). ** adjusted p-value < 0.01. **** adjusted p-value < 0.0001 (one-way ANOVA).
